# Thermoneutrality induces skeletal muscle myopathy via brown adipose tissue in an IRF4- and myostatin-dependent manner

**DOI:** 10.1101/274365

**Authors:** Xingxing Kong, Peng Zhou, Ting Yao, Lawrence Kazak, Danielle Tenen, Anna Lyubetskaya, Brian A. Dawes, Linus Tsai, Barbara B. Kahn, Bruce M. Spiegelman, Tiemin Liu, Evan D. Rosen

**Affiliations:** Division of Endocrinology, Beth Israel Deaconess Medical Center, Harvard Medical School, Boston, MA 02215; Division of Pediatric Endocrinology, Department of Pediatrics, David Geffen School of Medicine at UCLA, Los Angeles, CA 90095; State Key Laboratory of Genetic Engineering, Fudan University; Department of Endocrinology and Metabolism, Zhongshan Hospital, Fudan University; School of Life Sciences, Fudan University; Dana-Farber Cancer Institute, Harvard Medical School, Boston, MA 02215; Broad Institute, Cambridge, MA 02142

## Abstract

Skeletal muscle and brown adipose tissue (BAT) share a common lineage and have been functionally linked, as exercise increases browning through the actions of various myokines. It is unknown, however, whether BAT can affect skeletal muscle function. Our prior work has shown that loss of the transcription factor IRF4 in BAT (BATI4KO) reduces adaptive thermogenesis. Here, we note that these mice also have reduced exercise capacity relative to wild-type littermates, associated with diminished mitochondrial function, ribosomal protein synthesis, and mTOR signaling in muscle, in addition to the signature ultrastructural abnormalities of tubular aggregate myopathy. Within brown adipose tissue, loss of IRF4 caused the induction of a myogenic gene expression signature, which includes an increase in the secreted factor myostatin, a known inhibitor of muscle function. Reduction of myostatin activity by the injection of neutralizing antibodies or soluble ActRIIB receptor rescued the exercise capacity of BATI4KO mice. Additionally, overexpression of IRF4 in brown adipocytes reduced serum myostatin and increased mitochondrial function and exercise capacity in muscle. A physiological role for this system is suggested by the observation that mice housed at thermoneutrality show lower exercise capacity with increased serum myostatin; both of these abnormalities are corrected by surgical removal of BAT. Collectively, our data point to an unsuspected level of BAT-muscle cross-talk driven by IRF4 and myostatin.

**Highlights:**

1. Mice lacking IRF4 in BAT have a decrease in exercise capacity, accompanied by histological, ultrastructural, signaling, gene expression, and bioenergetic evidence of myopathy in white vastus.
2. Loss of IRF4 promotes the expression of a myogenic signature in BAT, including the myokine myostatin.
3. Neutralization of serum myostatin rescues the ability of BATI4KO mice to exercise normally, while overexpression of IRF4 in BAT allows mice to run better than wild-type counterparts.
4. Thermoneutrality reduces the level of IRF4 in BAT of WT mice, resulting in a myopathic phenotype that can be reversed by surgical excision of BAT.

## INTRODUCTION

Mammalian brown adipose tissue (BAT) is made up of specialized adipocytes that express uncoupling protein 1 (UCP1), which dissipates the mitochondrial proton gradient, forcing increased flux through the electron transport chain and subsequent heat generation. BAT activation, either by cold exposure or through the actions of β-adrenergic agonists, is associated with reduced body weight and improved glucose and lipid homeostasis in rodents and humans (Kajimura et al., 2015; Schrauwen and van Marken Lichtenbelt, 2016; Vosselman et al., 2013). These actions of BAT on systemic physiology are believed to be cell autonomous, with increased UCP1 expression leading to enhanced substrate consumption and thermogenesis. Additional UCP1-independent thermogenic pathways have recently been described in BAT, but these are also cell autonomous in nature (Ikeda et al., 2017; Kazak et al., 2015; Kazak et al., 2017).

Conversely, most of the actions of white adipose tissue on systemic physiology are believed to be mediated by the release of adipokines: small molecules, lipids, and proteins secreted by adipocytes that play an important role on appetite, satiety, insulin action, blood pressure, coagulation, and other homeostatic processes (Rosen and Spiegelman, 2014). Few examples of such factors have been shown to be involved in BAT-related biology, although this may be beginning to change. For example, neuregulin 4, is released from brown adipocytes and plays a role in regulating hepatic lipogenesis (Wang et al., 2014). It is likely that many such “BATokines” remain to be discovered.

We originally identified the transcription factor IRF4 as a regulator of adipogenesis after a DNase hypersensitivity screen revealed an interferon-stimulated response element (ISRE) as an over-represented motif in chromatin regions that become progressively accessible during differentiation (Eguchi et al., 2008). Subsequent work showed that IRF4 is strongly induced by fasting in adipocytes, upon which it simultaneously induces the expression of the enzymatic machinery of lipolysis and represses the corresponding genes that promote lipogenesis (Eguchi et al., 2011). More recently, IRF4 was identified as the transcriptional partner of the co-factor PGC-1α and a critical regulator of mitochondrial biogenesis and thermogenesis in BAT. Mice lacking IRF4 in BAT (BATI4KO) are sensitive to cold exposure and have increased adiposity and insulin resistance on a high fat diet, while mice overexpressing IRF4 in BAT (BATI4OE) have the opposite phenotype (Kong et al., 2014). The BATI4KO and BATI4OE are thus excellent models of deficient and hyper-efficient BAT function, generally.

One of the many features of brown fat biology that have gained attention recently is the propensity for exercise to promote browning of white adipose tissue, inducing the formation of thermogenic beige adipocytes with many of the same attributes as interscapular brown fat. Although it is still unclear what purpose this serves in the exercising animal, it is well understood that this occurs through the elaboration of myokines like irisin/FNDC5, Metrnl, and β-aminoisobutyric acid (Bostrom et al., 2012; Rao et al., 2014; Roberts et al., 2014). Intrigued by these data, we asked whether the skeletal muscle **→** BAT axis could also communicate in the reverse direction; that is, could skeletal muscle activity be under control of factors from BAT?

## RESULTS

### Mice lacking IRF4 in BAT have reduced exercise capacity

Male, chow-fed BATI4KO mice subjected to a low intensity treadmill regimen demonstrated diminished exercise capacity (**Fig. 1A, Figs. S1A, S1B**). Similar results were noted using a high intensity regimen, as well as free wheel running (**Fig. 1B, Figs. S1C-E**). These mice have targeted ablation of IRF4 in both interscapular BAT and beige adipocytes that exist within white depots. To distinguish which type of thermogenic adipocyte accounts for this effect on running ability we studied adipose-specific Prdm16-deficient mice, which have normal interscapular BAT but are deficient in beige adipocytes (Cohen et al., 2014). These mice exercise normally, suggesting that beige adipocytes are not major drivers of the effect on skeletal muscle (**Fig. 1C**). We also tested mice lacking UCP1 to determine if the thermogenic actions of BAT are required for the effect to take place, but again, we saw no difference in exercise capacity in this model (**Fig. 1D**). Importantly, we confirmed that our strategy for deleting IRF4 in BAT was not targeting *Irf4* in muscle inappropriately (**Fig. S1F**).

**Figure 1:**
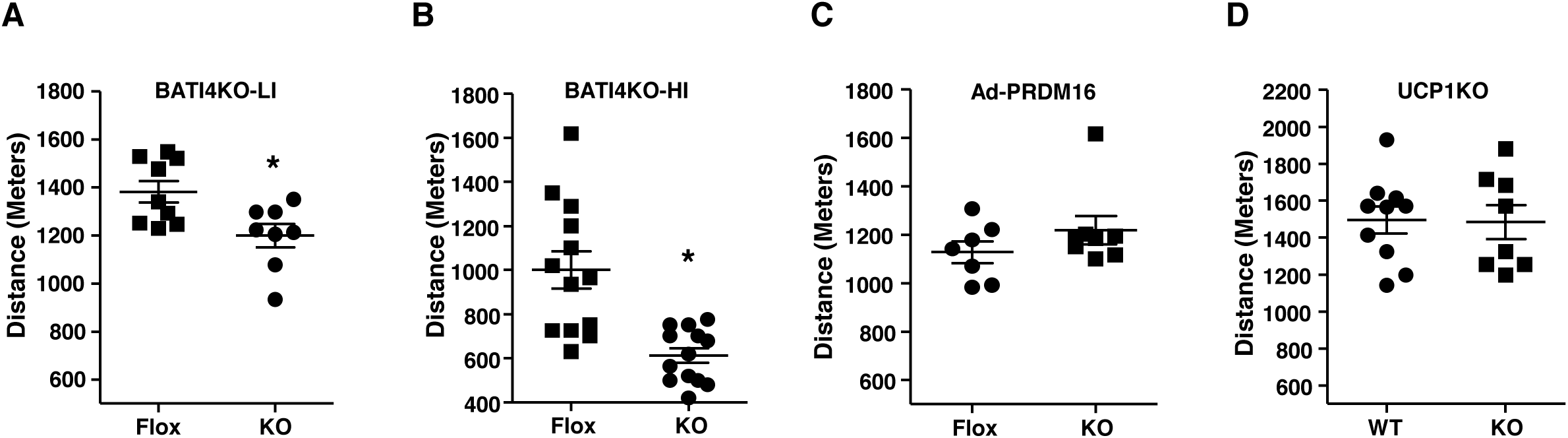
Mice lacking IRF4 in BAT have reduced exercise capacity. BATI4KO mice showed lower exercise capacity with both a Low Intensity (LI) **(A)** and High Intensity (HI) **(B)** regimen. The low intensity exercise regimen was performed in adipose-specific PRDM16 KO (**C**) and UCP1KO male mice (**D**). \**p* < 0.05.

### Mice lacking IRF4 in BAT have abnormalities of white vastus muscle

The skeletal muscles of BATI4KO mice appear grossly normal, with a normal distribution of Type I and Type II fibers (**Fig. S2A**). There were, however, signs of myopathy in at least some muscles. Specifically, in the white vastus lateralis there is nuclear relocation from the periphery to the center of the myofibers after running (**Fig. 2A**), a hallmark of dysfunctional muscle with or without degeneration/regeneration (Roman and Gomes, 2017). This was not seen in control mice, or in the soleus of BATI4KO mice (**Fig. S2B**). Even more strikingly, electron microscopy revealed the presence of tubular aggregates within the white vastus of BATI4KO mice (**Fig. 2B**), a pathological change associated with a variety of myotonic disorders found almost exclusively in Type II myocytes (Agbulut et al., 2000; Morgan-Hughes, 1998). Tubular aggregates are believed to derive from expansion of the sarcoplasmic reticulum (SR) (Funk et al., 2013); consistent with this we found greatly enhanced expression of the gene encoding the SR protein Atp2a2 (also called SERCA2) in affected muscles (**Fig. 2C**).

**Figure 2:**
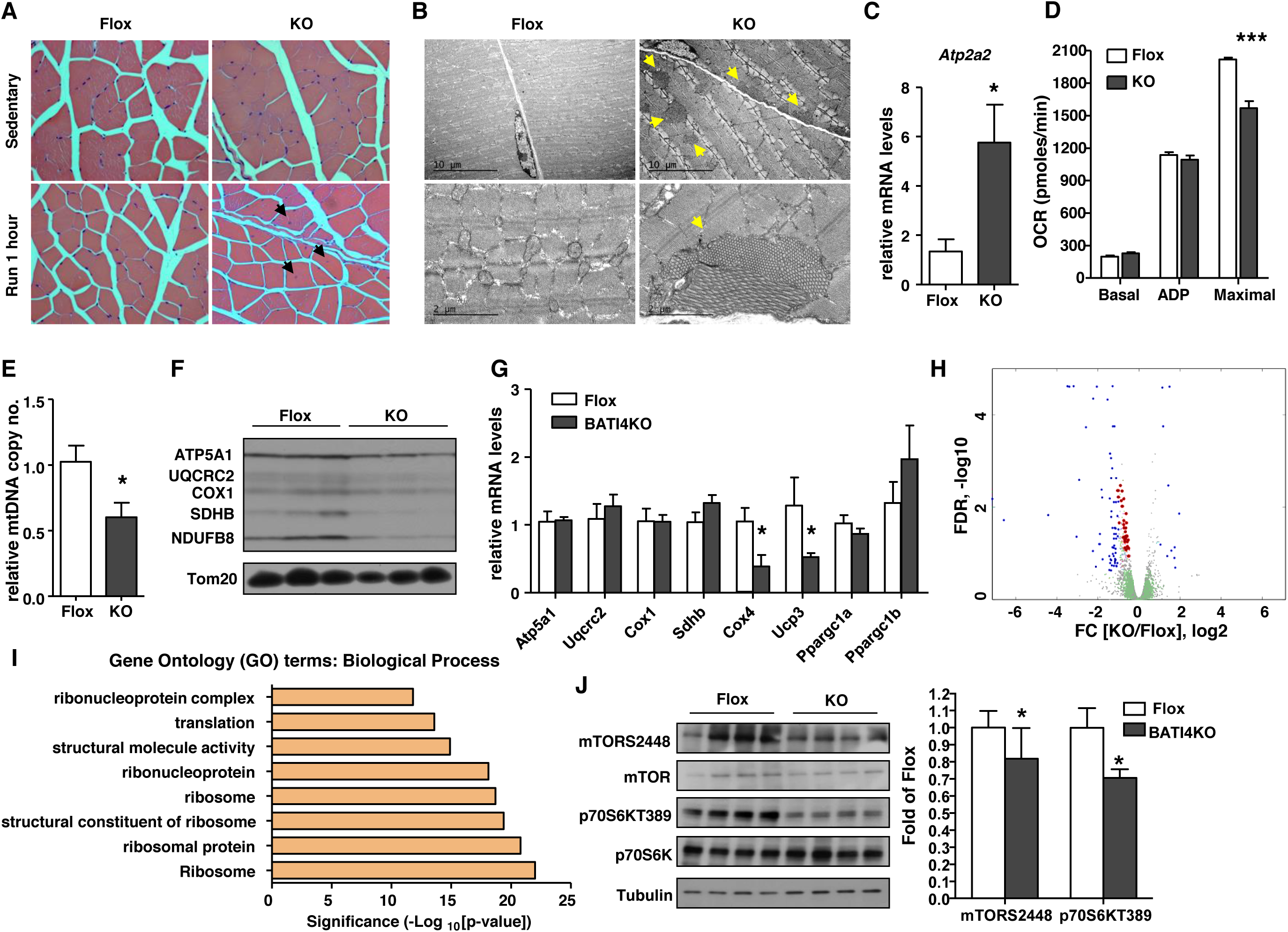
BATI4KO mice exhibit myopathy. **(A)** H&E staining of vastus lateralis of BATI4KO and control mice (X40). Black arrows indicate nuclei that have migrated to the center of the myofiber after 1 hour of running. (**B)** Tubular aggregates (yellow arrows) in the vastus lateralis of male BATI4KO mice. **(C)** *Atp2a2* expression in the vastus lateralis of male BATI4KO and control mice. Results are expressed as mean ± SEM (n=5-6, \**p* <0.05). (**D)** Measurement of oxygen consumption rate (OCR) in isolated mitochondria from vastus of male BATI4KO and control mice in the sedentary state. (n=7 mice per group, \*\*\**p*<0.0001). (**E)** Mitochondrial DNA content in the vastus lateralis of male BATI4KO and control mice. Results are expressed as mean ± SEM (n=4-5, \**p*<0.05). (**F)** Western blot analysis of isolated muscle mitochondria from BATI4KO and control mice. (**G)** Q-PCR analysis of mitochondrial gene expression in the vastus of male BATI4KO and control mice (n = 8-10, \**p*<0.05). (**H)** Volcano plot of vastus RNA-seq. Blue and red dots are significantly different between BATI4KO and flox mice; red dots represent ribosomal subunit genes. **(I)** GO analysis (biological process) of significantly different genes from RNA-seq analysis. **(J)** Western blot analysis of mTOR signaling in vastus of male BATI4KO vs. control mice. Activity was quantified using a phosphorimager. (n=4-7, \**p* < 0.05 vs. Flox mice.)

To ascertain whether reduced running ability was associated with a metabolic defect in affected muscles, we extracted mitochondria from the vastus lateralis of BATI4KO and control mice and measured the oxygen consumption rate (OCR). Basal (State 4) and ADP-stimulated respiration were unaltered, however, maximal OCR was much lower in the mitochondria from BATI4KO mice compared to control mice (**Fig. 2D**), suggesting decreased electron flux through the mitochondrial respiratory chain. This was associated with reduced mitochondrial DNA content and electron transport protein expression in purified vastus mitochondria (**Figs. 2E, 2F**), without any changes in soleus (**Figs. S2C, S2D**). We anticipated that these alterations in mitochondrial protein expression would reflect reduced mRNA expression of these genes, but this was not the case. RNA-seq analysis from white vastus of BATI4KO mice revealed diminished expression of a few mitochondrial genes (e.g. *Cox4, Ucp3*), but the majority of mitochondrial proteins displayed unchanged levels of mRNA (**Figs. 2G, 2H**). In contrast, gene set analysis suggested that ribosomal subunit genes were broadly down-regulated (**Figs. 2H, 2I, S2E, S2F, Suppl Table 1**).

Ribosomal subunit synthesis is under the control of mTOR signaling (Mayer and Grummt, 2006), leading us to consider whether mTOR signaling might be reduced in muscle from BATI4KO mice. In support of this idea, TORC1 (raptor) ablation in skeletal muscle causes a form of muscular dystrophy with features reminiscent of BATI4KO mice, such as reduced oxidative capacity and centrally located nuclei (Bentzinger et al., 2008). Indeed, we found significantly reduced phosphorylation levels of the mTOR target p70S6K (T389) and a trend towards reduced mTOR phosphorylation (S2448) in the vastus of BATI4KO mice (**Fig. 2J**).

### Loss of IRF4 induces a myogenic gene expression signature in BAT, including the TGFβ-family member myostatin

Based on these data, we speculated that a myopathic factor was being secreted from the BAT of BATI4KO mice. We therefore performed RNA-seq on interscapular BAT from BATI4KO and control mice, which revealed 530 differentially regulated genes corresponding to |log_2_ fold-change| ≥ 0.5 and FDR ≤ 0.25; of the 415 up-regulated genes a huge number were related to myocyte differentiation and muscle function (**Figs. 3A, 3B, Fig. S3A, Suppl Table 2**).

**Figure 3:**
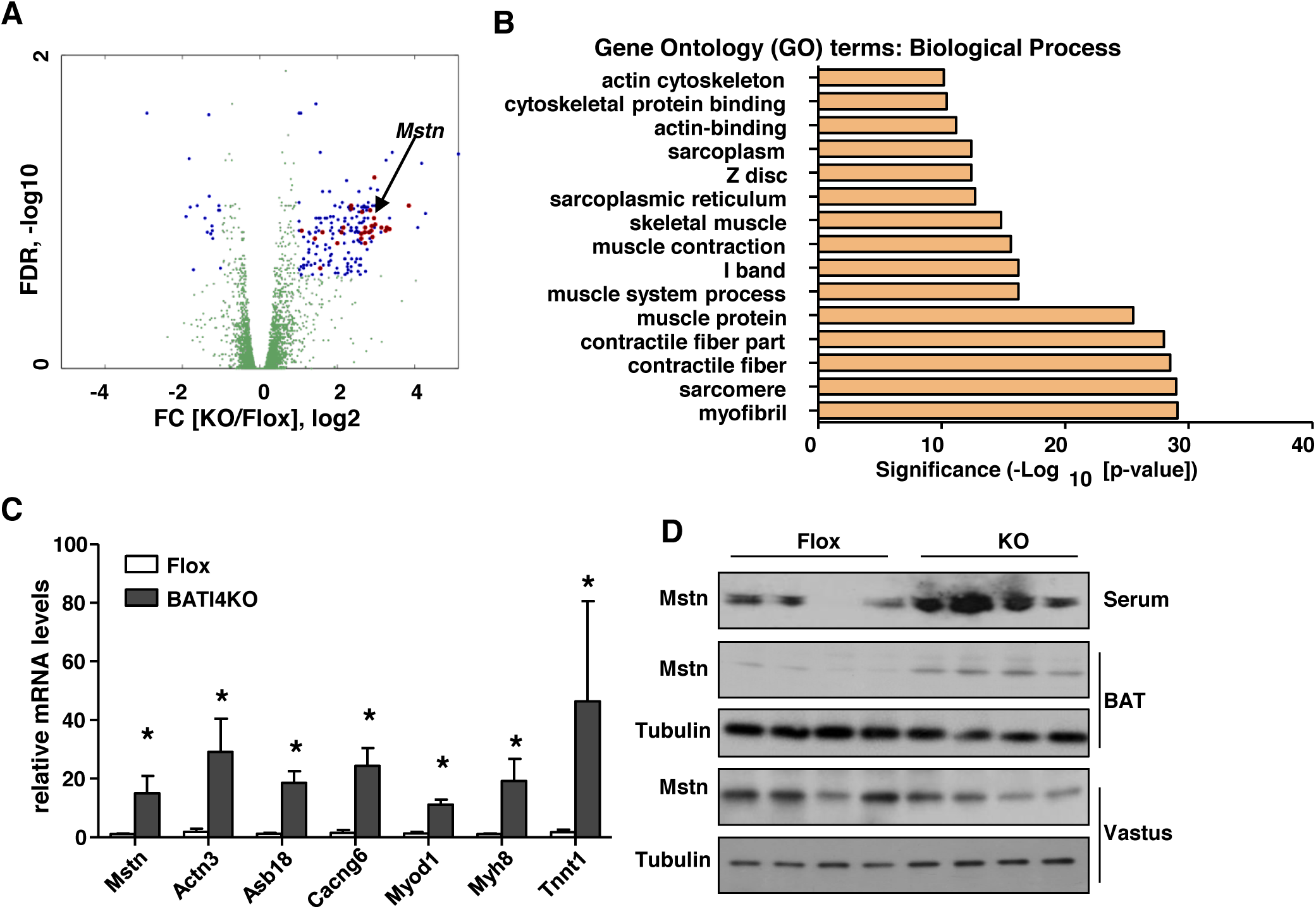
Loss of IRF4 induces a myogenic gene expression signature in BAT, including the TGFβ-family member myostatin. (**A)** Volcano plot of BAT RNA-seq. Blue and red dots represent significantly different genes between BATI4KO and flox groups. Red dots represent muscle-related transcripts. (**B)** GO analysis (biological process) of significantly different genes from RNA-seq analysis. (**C)** Q-PCR analysis of selected muscle-related genes in BAT of male BATI4KO vs. control mice (n=5-6, \**p*<0.05). (**D**) Tissue and serum myostatin levels in BATI4KO vs. control mice.

Interestingly, one of the strongest up-regulated genes was *Mstn*, encoding the secreted factor myostatin (**Figs. 3A 3C**), which is known to repress skeletal muscle hypertrophy in part via inhibition of mTOR signaling (Trendelenburg et al., 2009). Levels of *Mstn* protein were elevated in the BAT of BATI4KO mice, but not in muscle (**Fig. 3D, Fig. S3B**). Within BAT, the elevated myostatin was found in isolated adipocytes (**Fig. S3C**). Importantly, myostatin was also elevated in the serum of BATI4KO mice (**Fig. 3D**).

### Myostatin mediates the effect of BAT IRF4 loss on exercise capacity

To test whether myostatin could be the myopathic factor affecting BATI4KO mice, we first employed a gain-of-function approach. A single injection of myostatin reduced exercise capacity in wild-type mice, and also induced a similar gene expression profile in the white vastus (**Fig. 4A, Figs. S4A, 4B**). We next employed two distinct loss-of-function strategies to assess the effect of blocking myostatin on exercise capacity in BATI4KO mice. In the first, we utilized single dose administration of soluble activin receptor-Fc complex (sActRIIB-Fc)(Cadena et al., 2010), which successfully rescued running ability in BATI4KO mice (**Fig. 4B**). There was no effect in control mice, and the effect washed out after ten days, suggesting that the effect was not due to a hypertrophic effect on the muscles of treated mice. Because sActRIIB-Fc can bind other TGFβ-family members in addition to myostatin, we next treated mice with a specific neutralizing anti-myostatin antibody (α-Mstn). A single injection of α-Mstn completely restored the ability of BATI4KO mice to run normally by 60 hrs (**Fig. 4C**). Gene expression patterns in vastus were also normalized by α-Mstn (**Fig. 4D**). Again, re-testing mice ten days after injection demonstrated that the effect is transient, and therefore not due to reprogramming or hypertrophy of muscle (**Fig. 4C**).

**Figure 4:**
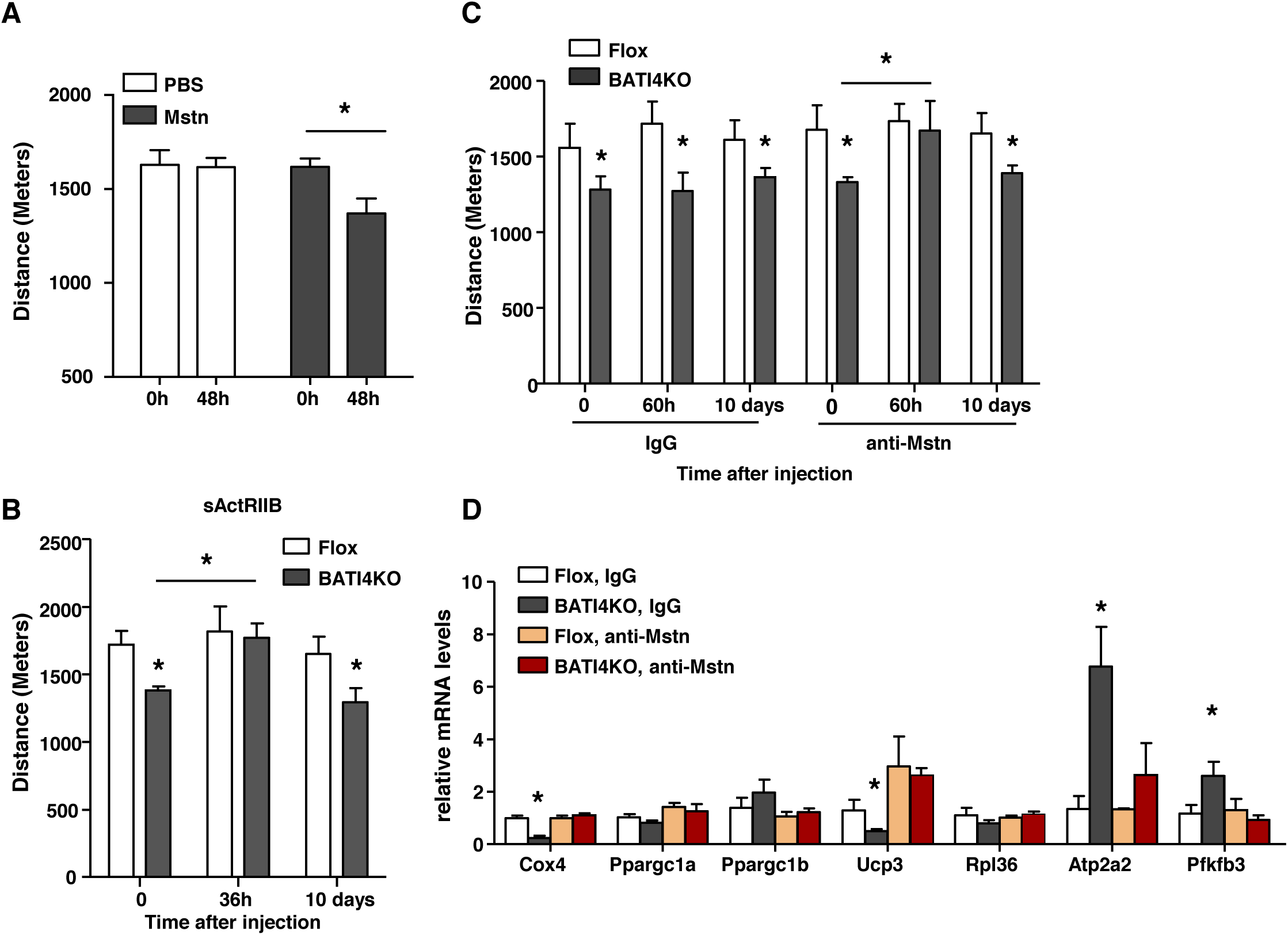
Myostatin mediates the effect of BAT IRF4 loss on exercise capacity. (**A**) Low intensity exercise regimen was performed in mice following a single injection of myostatin. (n=12-16, \**p* < 0.05.). (**B**) Exercise capacity determined in BATI4KO and control littermates before, 36 hours or 10 days following intraperitoneal injection of soluble ActRIIB (sActRIIB) with 10 mg/kg per mouse. (n=6-7, \**p*<0.05). **(C)** Exercise capacity was measured in BATI4KO and control littermates before, 60 hours or 10 days following a single intraperitoneal injection of myostatin neutralizing antibodies or isotype control. (n=6-7, \**p*<0.05). (**D)** Q-PCR analysis of gene expression in vastus lateralis from sedentary mice treated as in (**C**), but samples were harvested before and 60 hours after injection (n=5, \**p*<0.05).

### Mice overexpressing IRF4 in BAT run better than wild-type mice, and have reduced serum myostatin

We next tested whether overexpression of IRF4 in BAT (BATI4OE) could improve running ability in mice. BATI4OE mice show enhanced thermogenesis, and are lean on a high fat diet (Kong et al., 2014). On chow diet, these mice exhibit higher exercise capacity (**Fig. 5A**). No clear morphological differences were noted in the vastus lateralis of these animals relative to controls by either light or electron microscopy (**Figs. S5A-C**). However, ADP-stimulated and maximal mitochondrial respiration rates were increased in BATI4OE mice, as was mtDNA content (**Figs. 5B, 5C**) in vastus lateralis, but not soleus (**Fig. S5D**). Furthermore, serum myostatin levels were decreased in BATI4OE mice (**Fig. 5D**), and levels of *Mstn* mRNA and protein were decreased in the BAT, but not the muscle, of BATI4OE mice (**Fig. 5D, Fig. S5E**). Consistent with this, mTOR signaling in vastus from BATI4OE mice was enhanced relative to littermate controls (**Fig. 5E**).

**Figure 5:**
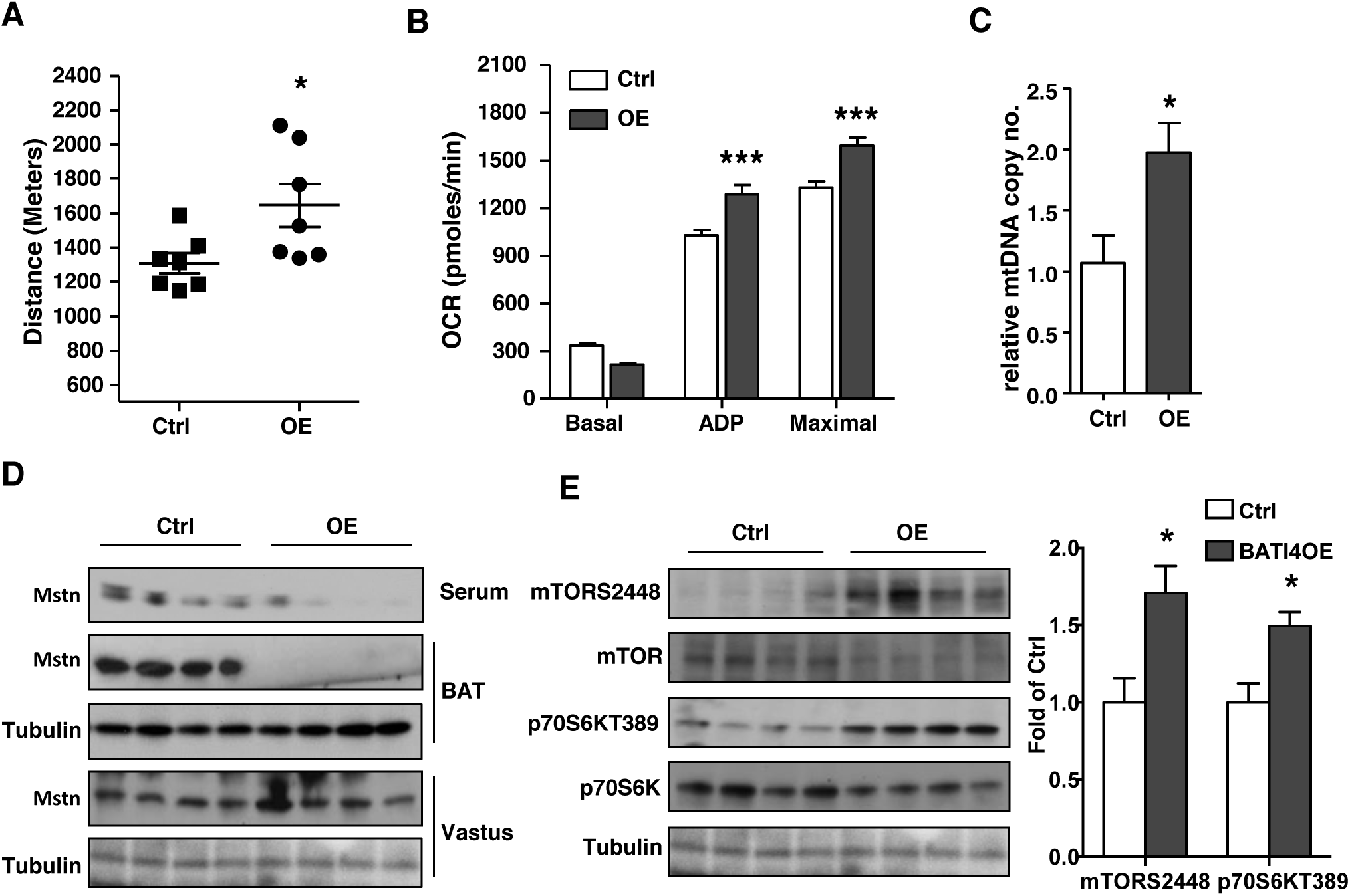
Mice overexpressing IRF4 in BAT run better than wild-type mice, and have reduced serum myostatin. (**A**) Exercise capacity in male BATI4OE mice (n=7, \**p*<0.05). (**B**) Continuous measurement of oxygen consumption rate (OCR) in isolated mitochondria from vastus of BATI4OE and control mice in the sedentary state. (**C**) Mitochondrial DNA content in the vastus of male BATI4OE and control mice. Results are expressed as mean ± SEM (n=4-5, \**p*<0.05). (**D**) Serum and tissue myostatin level in BATI4OE vs. control mice. (**E)** Western blot analysis of mTOR signaling pathway in vastus lateralis of BATI4OE and control mice. Activity was quantified using a phosphorimager. (n=4, \**p* < 0.05 vs. control mice.)

### Thermoneutrality induces expression of myostatin in BAT and reduces exercise capacity

In order to put these findings into physiological context, we placed wild-type mice at thermoneutrality (30^º^C), a condition that profoundly reduces *Irf4* expression (**Fig. S6A**). As predicted, warming resulted in a phenotype that is remarkably similar to BATI4KO; these mice show reduced exercise capacity associated with elevated tissue and serum myostatin concentrations. They also exhibit diminished mitochondrial protein expression and mTOR signaling in white vastus (**Figs. 6A, 6B, and Figs. S6B, C**). In addition, exposure to thermoneutrality induces a gene expression profile in vastus that is remarkably similar to that of BATI4KO mice (**Figs. S6D, E**). To prove that thermoneutrality exerts these effects via BAT, we surgically removed interscapular BAT (iBATx) immediately prior to placement at 30^º^C. After one week of recovery (**Fig. S6F**), iBATx mice showed improved exercise capacity relative to sham-operated controls, with decreased myostatin levels in serum (**Figs. 6C, 6D**). Vastus mitochondrial protein levels, mTOR signaling, and mitochondrial and ribosomal gene expression were all normalized in iBATx mice relative to sham-operated controls (**Figs. 6E, and Figs. S6G-I**).

**Figure 6:**
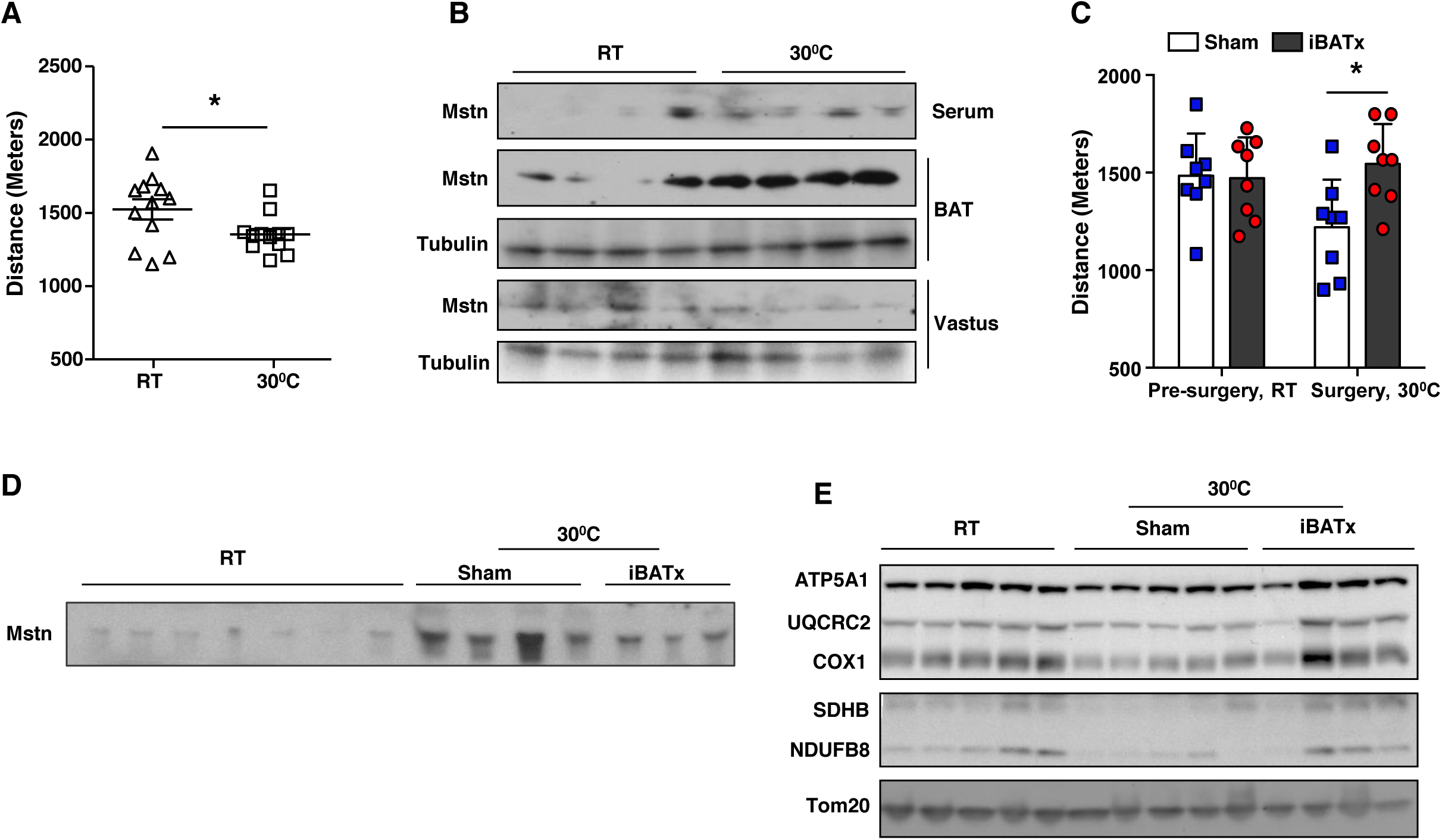
Thermoneutrality induces expression of myostatin in BAT and reduces exercise capacity. **(A)** Exercise capacity was measured in male wild-type C57Bl/6J mice after exposure to 30°C for 7 days, (n=10-12, \**p*<0.05). (**B)** BAT, vastus, and serum levels of myostatin in mice housed at thermoneutrality vs. RT. (**C**) Exercise capacity in male iBATx mice (n=8, \**p*<0.05). (**D**) Serum myostatin level in mice before and after iBATx and exposure to 30°C. (**E)** Western blot analysis of isolated muscle mitochondria from mice before and after iBATx and exposure to 30°C.

## DISCUSSION

BAT has long been thought to have one major function—generation of heat, which occurs when the mitochondrial proton gradient is dissipated by uncoupling protein 1. This simplistic notion has given way of late to a more sophisticated and nuanced perspective that concerns at least three different areas of BAT biology. First, a critical distinction has been made between interscapular brown adipose tissue and the “brown-like” cells that appear within nominally white depots during cold exposure. These latter cells, now called beige (or BRITE) adipocytes, are thermogenic cells that express UCP1 like interscapular brown adipocytes, but which derive from a different lineage and possess an overlapping but distinct gene signature (Harms and Seale, 2013). Second, the observation that animals lacking UCP1 can tolerate cold (Hofmann et al., 2001; Liu et al., 2003) paved the way for the elucidation of UCP1-independent mechanisms of thermogenesis. These include a futile cycle between creatine and creatine kinase that was originally thought to occur only in beige adipocytes, but is now appreciated in brown cells as well (Kazak et al., 2015; Kazak et al., 2017). More recently a SERCA2-based calcium cycle was shown to mediate heat generation in beige fat (Ikeda et al., 2017), and other UCP1-independent pathways for heat generation are under investigation. Third, the observation that ablation of BAT causes more profound metabolic consequences than knocking out UCP1 (Enerback et al., 1997; Lowell et al., 1993) led to the proposition that BAT does something beyond simple heat generation. This idea was bolstered by the discovery of adipokines, secreted products of adipocytes that coordinate various aspects of energy homeostasis and metabolic physiology (Rosen and Spiegelman, 2014). In addition to a panoply of adipokines that are shared with white adipocytes, brown adipocytes are likely to secrete some factors that are unique to thermogenic cells. One such example is neuregulin 4, an EGF-like protein secreted by BAT that represses hepatic lipogenesis (Wang et al., 2014). Our discovery of a BAT-muscle axis that regulates exercise capacity is therefore in line with this burgeoning sense of what thermogenic adipocytes are, and what they can do.

A little more than a decade ago, similarities were noted in the transcriptional profiles of skeletal muscle and brown adipose precursor cells (Timmons et al., 2007). Additional studies using lineage tracing techniques suggested that there might be a common Engrailed+ precursor that gives rise to both skeletal muscle and brown fat from a common central dermomyotome (Atit et al., 2006). Finally, brown adipocytes and skeletal muscle were shown to share a common Pax7^+^, Myf5^+^ lineage, and the transcription factors Ebf2, Ppar*γ*, and Prdm16 were identified as key determinants favoring adipogenesis over myogenesis (Rajakumari et al., 2013; Seale et al., 2008). Our data suggest that Irf4 may play an important role in this process as well. Of note, loss of Irf4 in BAT does not cause a full reprogramming to muscle; Irf4 null brown adipocytes look like wild-type cells at the light microscopic and ultrastructural level (Kong et al., 2014). They also continue to express most of the genes of wild-type brown adipocytes, albeit with some (e.g. *Ucp1, Ppargc1a*) at reduced levels (Kong et al., 2014). Instead, there is the appearance of a broad myogenic signature at low overall levels that probably results from actions at a few critical myogenic drivers, like MyoD1. Presumably, the full muscle phenotype is not elaborated because the key repressors of their expression, such as Ebf2, Ppar*γ*, and Prdm16, are still present.

Our BATI4KO data bring to mind recent work by the Brüning group, who found that optogenetic and chemogenetic activation of hypothalamic Agrp^+^ neurons can promote the expression of a myogenic signature in BAT almost exactly the same as we show here (Steculorum et al., 2016). This makes sense, as activation of Agrp^+^ neurons causes a sharp reduction in sympathetic activation of BAT (Krashes et al., 2011), which would be expected to cause a decrease in IRF4 levels in BAT (Kong et al., 2014), akin to the effect that we see following thermoneutrality. Interestingly, Brüning and colleagues also identified myostatin secretion from BAT, which they showed acts in an autocrine manner to reduce local insulin sensitivity; no effect on muscle insulin sensitivity was noted during a hyperinsulinemic-euglycemic clamp.

The effects of myostatin on muscle have been studied extensively using either massive overexpression or complete knockout, manipulations which cause significant atrophy or hypertrophy, respectively (Allen et al., 2011; Dschietzig, 2014). In contrast, in our models we see neither atrophy nor hypertrophy. We speculate that this is because the changes in serum myostatin are relatively modest compared to these prolonged exposures to pharmacological levels, or complete absence of, myostatin. We also speculate that this may explain why some muscles, like white vastus, are selectively affected.

Myostatin is believed to be produced primarily by muscle, and certainly the total amount of muscle in a mouse vastly exceeds the mass of BAT. Nevertheless, our data suggest that serum myostatin is disproportionately derived from BAT, at least under certain conditions. In the BATI4KO model, we see elevated serum myostatin that corresponds to increased myostatin mRNA and protein in BAT, but not muscle. Conversely, our analysis of the BATI4OE model shows reduction of myostatin in BAT and serum, but not muscle. Finally, at thermoneutrality, we see a large rise in serum myostatin by Western blotting that is largely reversed by BATectomy. The residual increment that remains after excision of BAT is likely due to the small pockets of subscapular BAT that are not removed in the operation. Taken together, these data provide significant evidence that murine serum myostatin derives from BAT, at least under certain environmental and genetic conditions. What portion of human serum myostatin derives from BAT, and under what conditions that may rise or fall, is unknown.

Is myostatin likely to be the only myopathic factor secreted by BAT at thermoneutrality, or in the presence of an IRF4 mutation? We show that a single dose of purified myostatin can cause the exercise phenotype. The studies in which myostatin is depleted by the soluble sActRIIB-Fc complex leading to complete restoration of exercise capacity leave open the possibility that related TGFβ family members, some of which can bind ActRIIB, may participate in the process. We did not see increased levels of such candidate factors in the IRF4 null BAT, but such a putative factor might not be regulated at the transcriptional level. The studies using a more specific antibody generated by Acceleron would seem to close that loophole; unfortunately the data indicating the specificity of this antibody are proprietary. We therefore conclude that myostatin is likely to be a dominant, if not the only, player in this axis.

Tubular aggregates were first described in the 1960s in human muscle biopsies from certain patients with weakness and myalgia (Funk et al., 2013; Schiaffino, 2012). Although there was some early speculation that they may represent deformed mitochondria, it is now well established that they derive from sarcoplasmic reticulum. They are seen preferentially in Type II muscle (especially Type IIb fibers, which comprise 95% of the white vastus in mice) and in males. There is a cluster of rare familial tubular aggregate myopathies in humans caused by mutations in the genes *STIM1* and *ORAI1*, among others yet to be identified (Bohm et al., 2013; Endo et al., 2015); these genes were not changed at the mRNA level in muscles from BATI4KO mice (not shown). In mice, tubular aggregates can arise in several pathological situations, including uremia and aging (Funk et al., 2013; Schiaffino, 2012). Of note, wild-type C57Bl/6 mice do not show evidence of tubular aggregate formation in vastus until they reach 14 months of age (Agbulut et al., 2000); our BATI4KO mice, which are on a C57Bl/6 background, were only 10 weeks of age when tested. There is only a single reference in the literature that we are aware of connecting myostatin to tubular aggregates. In that study, constitutive global knockout of myostatin was associated with increased muscle mass, in particular an increased number of Type IIb fibers, but with paradoxically reduced muscle force; tubular aggregate formation was noted in those fibers as well (Amthor et al., 2007). It is unclear how to put these results into context with our observations. Most likely tubular aggregate formation is simply a generic response to certain types of stress in murine Type IIb fibers.

To our knowledge, the effect of thermal stress on exercise has not been systematically studied in animal models, although there are reports suggesting that pre-warming has detrimental effects on exercise capacity in humans (Gregson et al., 2005; Levels et al., 2014). We speculate that an organism might want to reduce the chance of hyperthermia by restricting exercise efficiency at high temperatures; in such a paradigm brown fat might be an ideal effector as an organ responsible for integrating thermal signals. Intriguingly, there are data showing that cold exposure can be used as a “cross-training” modality in sparrows, such that cold exposed birds show enhanced muscle function and exercise performance; this is also associated with reduced serum myostatin (Zhang et al., 2015).

Finally, our models also resemble the situation in obesity, a condition with reduced functional BAT (van Marken Lichtenbelt et al., 2009), reduced IRF4 in BAT (Kong et al., 2014), elevated serum myostatin (Allen et al., 2011) and diminished ribosomal protein expression in skeletal muscle (Campbell et al., 2016). Conversely, bariatric surgery increases BAT activity (Vijgen et al., 2012), while also reducing serum myostatin (Park et al., 2006) and restoring ribosomal gene expression in muscle (Campbell et al., 2016).

In summary, we report here that brown adipose tissue can secrete significant quantities of myostatin into the blood in response to warming, a situation that is mimicked by deleting IRF4 in BAT. In both of these situations, myostatin causes reduced mTOR signaling in Type IIB muscle fibers, with a subsequent negative effect on translation. This seems to affect certain proteins preferentially, among them components of the mitochondrial electron transport chain. This in turn causes a bioenergetic defect in susceptible muscle, and a functional myopathy.

Taken together, our data provide evidence of an unsuspected level of inter-organ cross-talk between BAT and skeletal muscle, involving the transcription factor IRF4 and the secreted protein myostatin. Our results suggest that BAT dysfunction can promote a myopathic state, and that enhancing BAT function might be a useful strategy to improve exercise capacity.

## STAR METHODS

### Antibodies

Antibodies were purchased from Santa Cruz Biotechnology (Tom20, sc-11415), Abcam (PDH, AB110416; Mito profile, AB110413), and Cell Signaling Technology (mTOR, 2972; Phospho-mTOR (Ser2448), 2971; p70 S6 Kinase, 2708; Phospho-p70 S6 Kinase (Thr389), 9234; α/β-Tubulin, 2148).

### Animal Care

Mice were maintained under a 12 hr light/12hr dark cycle at constant temperature (23°C) with free access to food and water. BATI4KO, BATI4OE, and adipose-specific Prdm16 KO mice were generated as described(Cohen et al., 2014; Kong et al., 2014). All animal studies were approved by the Institutional Animal Care and Use Committee of the Beth Israel Deaconess Medical Center.

### Thermoneutrality stimulation

Male C57BL mice housed at 30°C were fed chow diet with free access to water and food for a week. Mice were maintained under a 12 hr light/12 hr dark cycle. After one week, mice were measured exercise capacity compared with those housed at room temperature. At the end of the studies, all animals were euthanized by cervical dislocation, trunk blood was collected and centrifuged (800 g, 20 min) to generate plasma and BAT and muscle tissues were removed, frozen in liquid nitrogen and stored at −80°C for later analyses.

### Surgical removal of interscapular BAT (iBATX)

iBATX animals were anaesthetized. A small longitudinal incision was made between the shoulder blades and the skin carefully opened. Sulzer’s vein draining the iBAT was located and tied off above the point of branching into the two iBAT lobes. The vein was then cut on the iBAT side of the tie and the two iBAT lobes were quickly and completely removed. The incision was closed and animals were allowed to recover. Control mice were given the same dose of anaesthesia, and had the skin opened and closed. All mice were treated with the pain-killer meloxicam (Zoopharm) before surgery. Mice were maintained under a 12 hr light/12 hr dark cycle at constant temperature (30^º^C) with free access to food and water and fed on a standard chow diet after surgery. Body weight was monitored every day until recovery(Rothwell and Stock, 1989).

### Measurement of exercise capacity

All mice were acclimated to the treadmill 4-5 days prior to the exercise test session. Before each session, food was removed 2 hours before exercise. Acclimation began at a low speed of 5 to 8 meters per minute (m/min) for a total of 10 minutes on Day 1, and was increased to 5 to 10 m/min for a total of 10 minutes on Day 2. Following this, the mice were allowed to rest for at least 2 days in their home cage. For the low intensity treadmill test, the treadmill began at a rate of 12 m/min for 40 minutes. After 40 minutes, the treadmill speed was increased at a rate of 1 m/min every 10 min for a total of 30 min, and then increased at the rate of 1m/min every 5 min until the mice were exhausted (mice spent more than 5 seconds on the electric shocker without resuming running). The high intensity treadmill test was conducted on an open field six-lane treadmill set at a 10% incline. Following a 5-min 0m/min acclimation period, the speed was raised to 6 m/min and increased by 2 m/min every 5 min to a maximal pace of 30m/min until exhaustion(DeBalsi et al., 2014). Individual experiments were replicated 2-4 times.

### Bioenergetics

Tissue respiration was performed using a Clark electrode (Strathkelvin Instruments). For mitochondrial bioenergetic analyses, vastus mitochondria were isolated in SHE buffer, pH 7.4 (250 mM sucrose, 5 mM Tris, pH 7.4, 1 mM EGTA) via differential centrifugation. Mitochondria were resuspended in respiration buffer (210 mM Mannitol, 70 mM Sucrose, 3 mM MgCl2, 5 mM KH2PO4, 20 mM Tris [pH 7.4], 0.1 mM EGTA, 0.1% BSA, 10 mM Pyruvate, 5 mM Malate). Oxygen consumption of vastus mitochondria (15 μg) was determined using an XF24e Extracellular Flux Analyzer (Seahorse Bioscience). ADP-stimulated respiration was triggered with 0.2 mM ADP. Maximal oxygen consumption was determined using FCCP (10 µM).

### Mitochondrial DNA copy number

Mitochondrial DNA (mtDNA) copy number was determined as a marker for mitochondrial density using quantitative RT-PCR as previously reported(Kong et al., 2010). Briefly, total DNA was isolated from the cells using DNeasy Blood & Tissue Kit (Qiagen, 69506) according to the manufacturer’s instructions. The mitochondrial DNA copy-number was calculated from the ratio of COX II (mitochondrial-encoded gene)/cyclophilin A (nuclear-encoded gene).

### RNA-seq library generation and analysis

Five replicates (each one an individual mouse) were prepared, sequenced, and analyzed separately. mRNA was purified from 70 ng of total RNA using the Ribo-Zero Magnetic Gold Kit (Illumina, catalog# MRZG126). Libraries were prepared using the TruSeq RNA Library Preparation Kit v2 (Illumina, catalog #RS-122-2001) according to the manufacturer’s protocol starting with the EPF step. Sequencing was performed on the Illumina HiSeq2500. RNA-seq data were aligned using TopHat2(Kim et al., 2013) to the mm9 mouse genome, and duplicates and low quality reads were removed by Picard (http://picard.sourceforge.net). Reads were assigned to transcripts, normalized and quantified using featureCounts(Liao et al., 2014) and EdgeR(Robinson et al., 2010). Low expressed genes (log2 CPM <1) were filtered out. Gene set enrichment analysis was performed using RDAVIDWebService(Fresno and Fernandez, 2013) and GOstats(Falcon and Gentleman, 2007).

### Myostatin tail vein injection

10 weeks old male C57bl mice were injected with myostatin (1ug/g BW) (Acceleron, Boston, MA, USA) or PBS through tail vein. 48 hours after injection, mice either underwent exercise capacity testing or were euthanized by cervical dislocation for sample collection; muscle tissues were removed, frozen in liquid nitrogen and stored at −80°C for later analyses. Individual experiments were replicated 2-4 times.

### Anti-myostatin blockade and sActRIIB injection

10-14 weeks old BATI4KO and control mice received intraperitoneal (IP) injections of anti-myostatin (αGDF8) or control IgG, or recombinant soluble ActRIIB (10 mg/kg) (Acceleron, Boston, MA, USA) in sterile TBS (1.0 mg/mL) (n=6-7). 60 hours (for anti-myostatin and IgG groups)/ 36hours (sActRIIB group) after the injection, mice were either measured exercise capacity or euthanized by cervical dislocation, trunk blood was collected and centrifuged (800 g, 20 min) to generate plasma and BAT and muscle tissues were removed, frozen in liquid nitrogen and stored at −80°C for later analyses. Individual experiments were replicated 2-4 times.

### Statistical Analysis

Unpaired two-tailed Student’s t-test and two-way ANOVA were used. *p*<0.05 was considered statistically significant.

### Data Availability Statement

All data used in this manuscript are presented in the figures and supplementary tables. Additional raw data or information can be obtained from the authors by request. RNA-seq datasets are available at GEO (GSE103736).

## ACKNOWLEDGEMENTS

The authors gratefully acknowledge the Electron Microscopy core of the BIDMC. John Shelton in the Molecular Pathology Core in the Division of Cardiology at UT Southwestern performed the ATPase staining. Scott Pearsall at Acceleron provided myostatin, Mstn antibody and the soluble ActRIIB receptor. We thank members of the Rosen and Spiegelman laboratories for helpful discussions and technical advice. This work was funded by R00 DK106550 to X.K., AHA 14SDG20370016 to T.L., NIH R01 DK31405 to B.M.S., NIH R37 DK43051 to B.B.K., and NIH R01 DK085171 and DK102170 to E.D.R.

## AUTHOR CONTRIBUTIONS

The experimental plan was designed by X.K., T.L. and E.D.R. X.K., T.L. P.Z., and L.K. performed experiments and analyzed data. D.T. generated RNA-seq data, which was analyzed by A.L. and L.T. X.K. and E.D.R. wrote the manuscript, incorporating edits and comments from B.B.K., B.M.S, and all other authors. E.D.R. is the senior and corresponding author.

---

**Figure S1:**
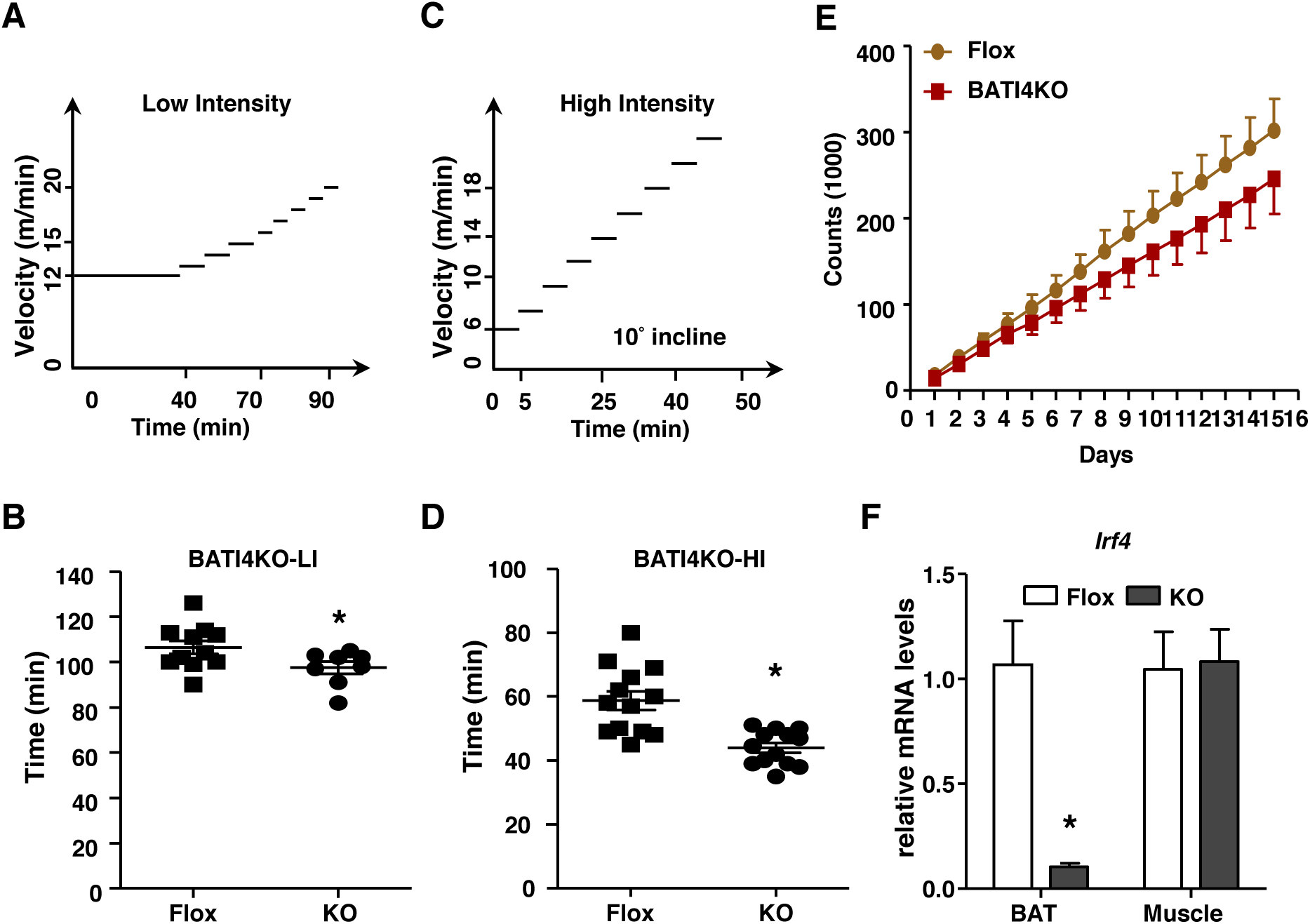
Reduced exercise capacity in BATI4KO mice. (**A**) Schematic of low intensity and (**C**) high intensity exercise regimens. Time run by BATI4KO and control mice on low intensity (**B**) and high intensity (**D**) protocols. (**E**) Cumulative free wheel running in BATI4KO and control mice (n=8 per group). **(F)** Q-PCR gene expression analysis of *Irf4* expression in BAT and vastus lateralis of BATI4KO mice. Results are expressed as mean ± SEM (n=5-6, \**p*<0.05).

**Figure S2:**
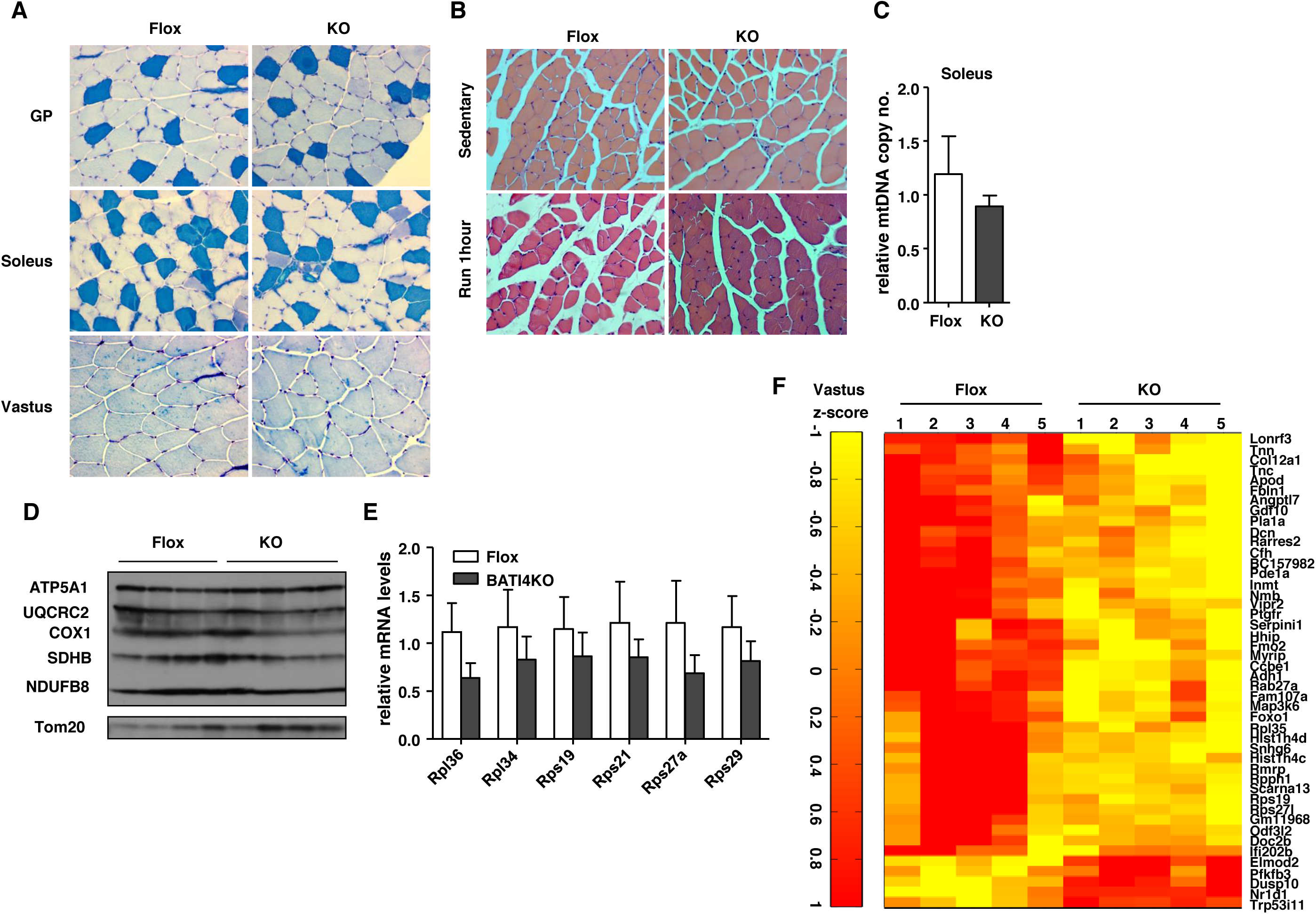
BATI4KO mice have altered skeletal muscle. **(A)** ATPase staining of muscle sections from BATI4KO and control mice (type I: dark blue; type IIa: white; type IIb: light blue). (**B**) H&E staining of soleus muscle from BATI4KO and control mice (40X). **(C)** Mitochondrial DNA number in soleus muscle of BATI4KO and control mice (n=5). **(D)** Western blot analysis of isolated muscle mitochondria from soleus of BATI4KO and control mice. **(E)** Q-PCR analysis of ribosomal protein gene mRNA expression in the vastus (n=8-10). **(F)** Heat map from RNA-seq of vastus lateralis from BATI4KO and control mice, |log2FC|>=1, FDR ≤ 0.05.

**Figure S3:**
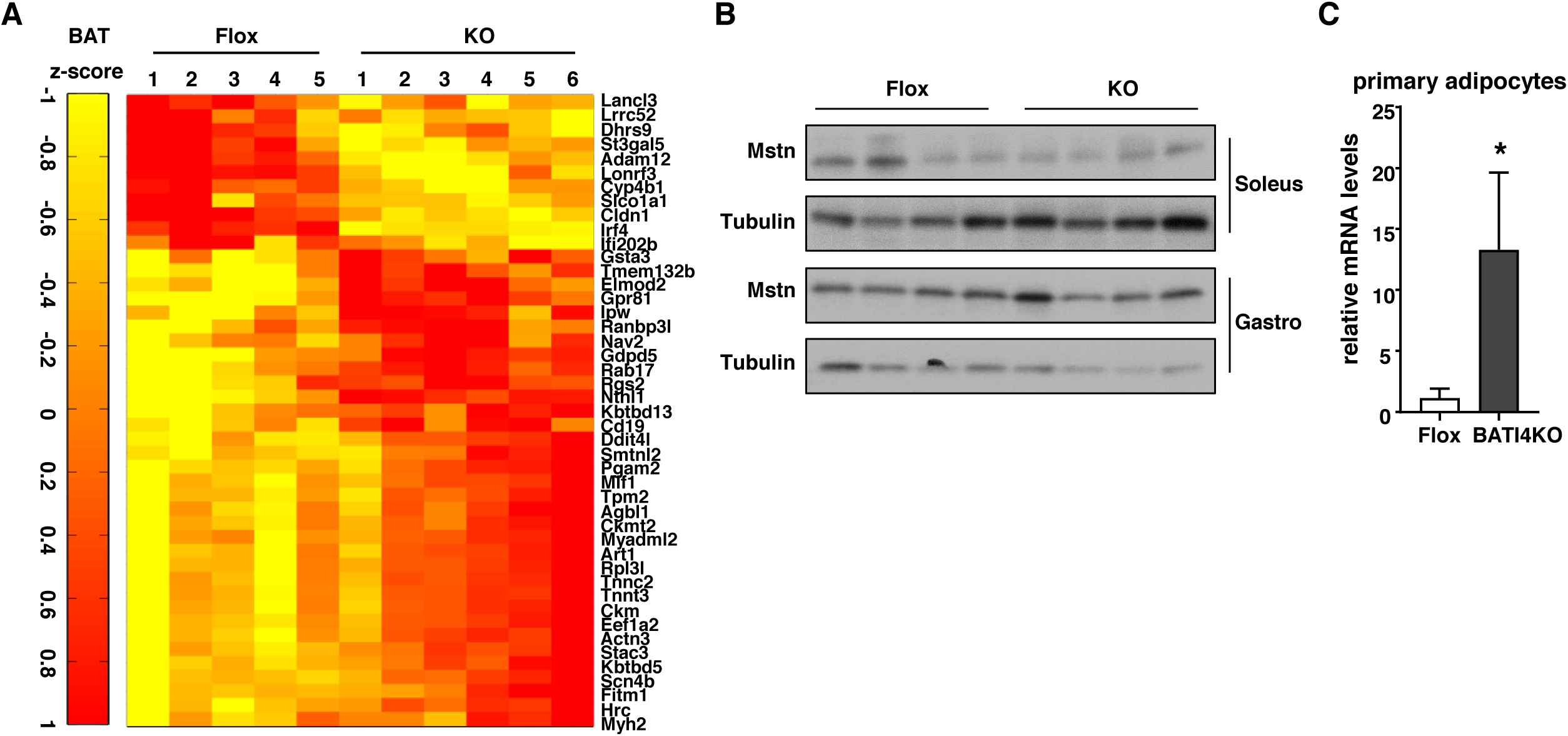
Direct effect of BAT via secretion of myostatin. (**A**) Heat map of differentially expressed genes in BAT of BATI4KO and control mice, |log2FC|>=1, FDR ≤ 0.05. (**B**) Western blot analysis of muscle myostatin levels. Gastro: gastrocnemius. (**C**) Elevated Mstn mRNA in primary brown adipocytes from BATI4KO mice. (n=5-6, \**p* < 0.05).

**Figure S4:**
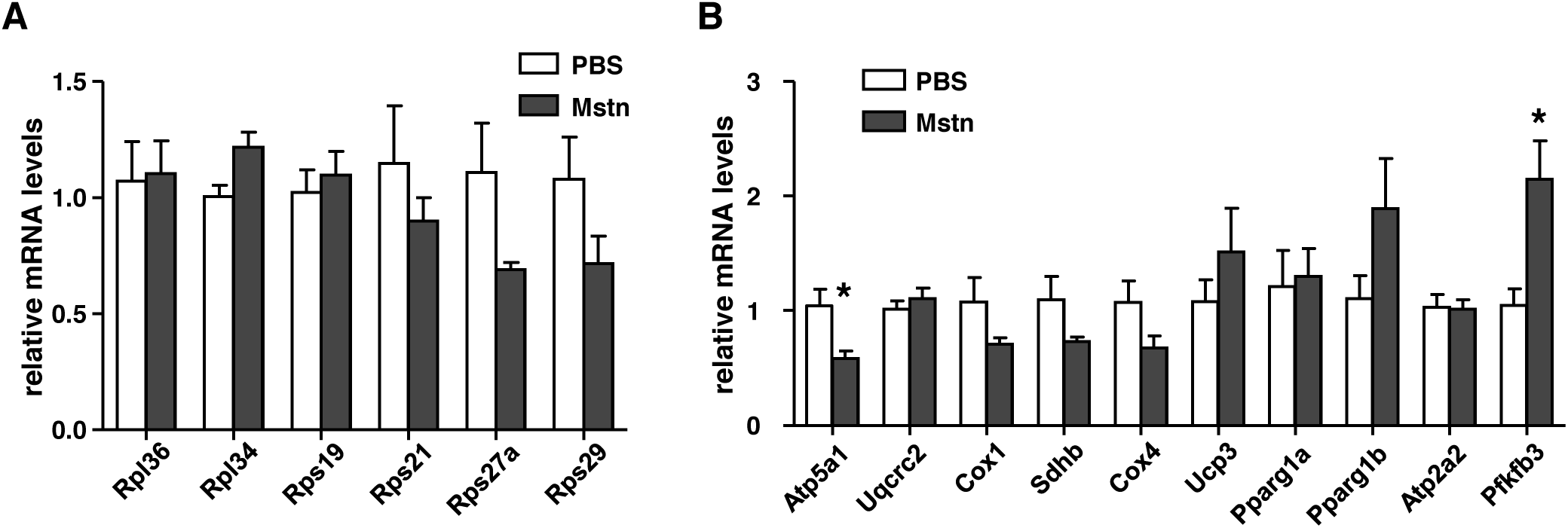
Myostatin mediates the effect of BAT IRF4 loss on exercise capacity. Q-PCR analysis of mitochondrial gene expression (**A**) and ribosomal protein gene mRNA expression (**B**) in the vastus from myostatin injection groups. (n=5-6, \**p* < 0.05).

**Figure S5:**
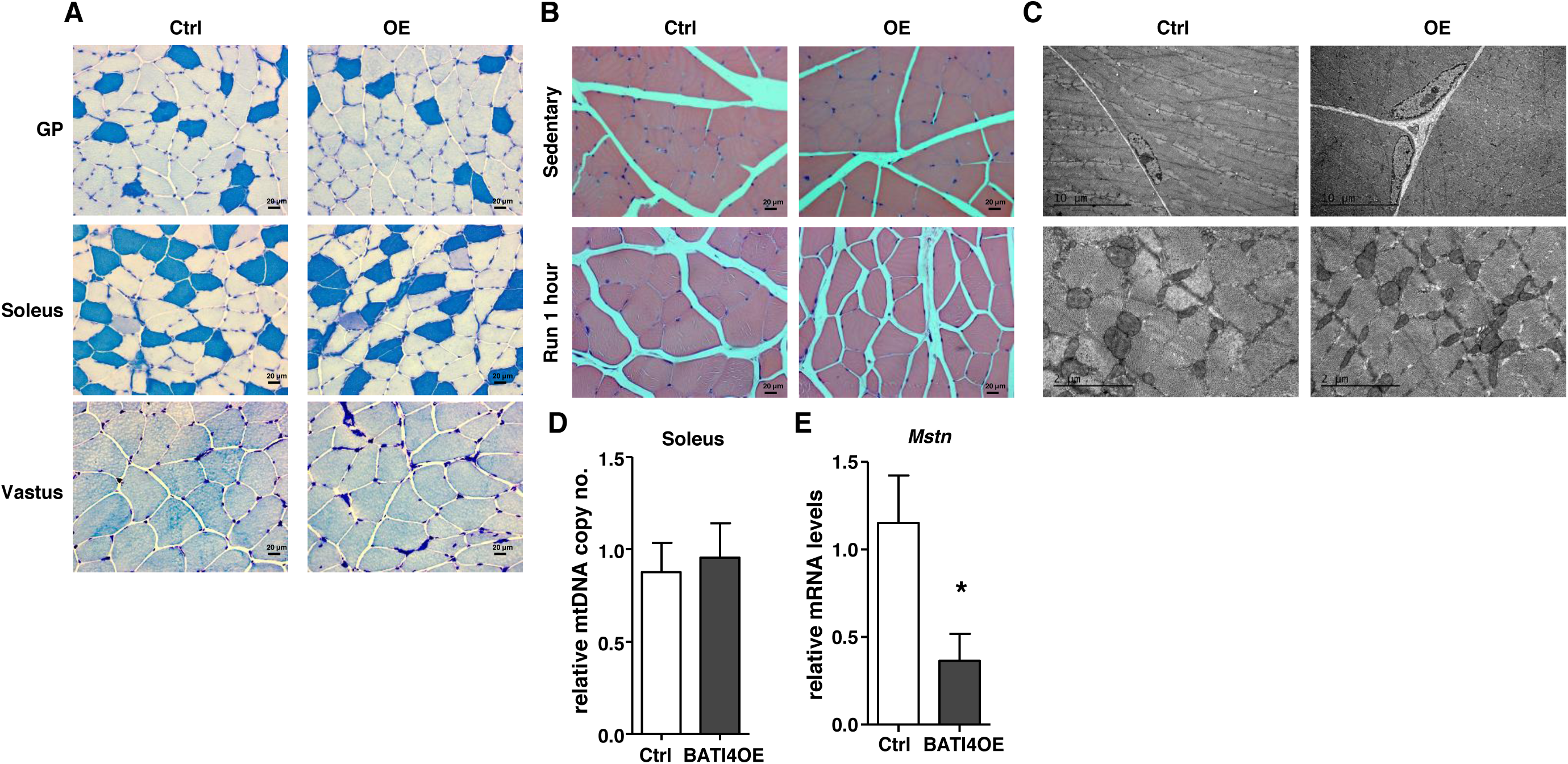
Increased exercise capacity following myostatin neutralization or IRF4 overexpression. **(A)** ATPase staining of muscle sections from BATI4OE and control mice (type I: dark blue; type IIa: white; type IIb: light blue). (**B)** H&E staining of vastus lateralis from BATI4OE vs. control mice (40X). (**C**) Electron micrograph of vastus lateralis from BATI4OE and control mice. (**D)** Mitochondrial DNA number in soleus of BATI4OE and control mice (n=4, 5). (**E)** Q-PCR analysis of *Mstn* mRNA expression in BAT (n=5-6, \**p*<0.05) of BATI4OE and control mice.

**Figure S6:**
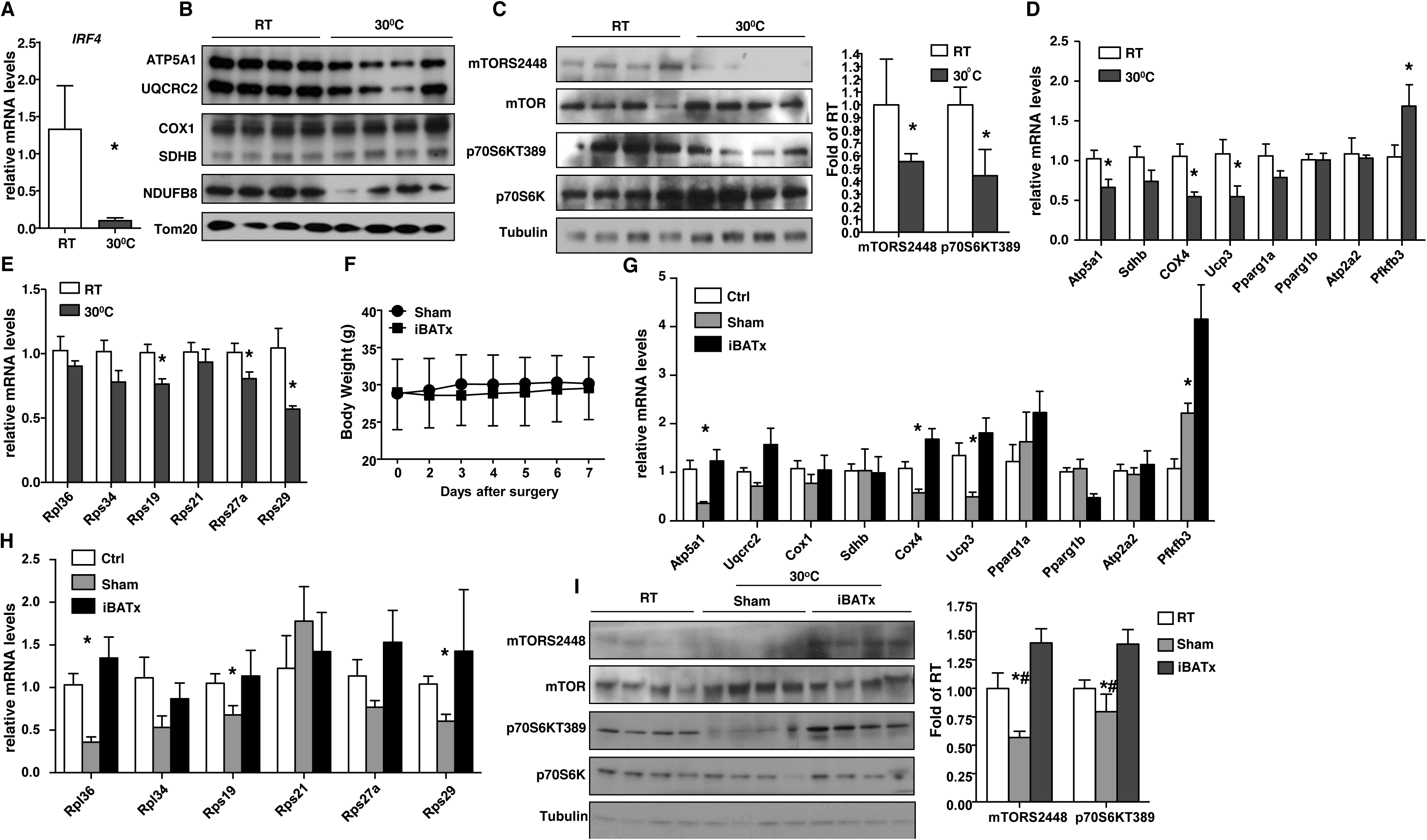
Warming causes myostatin elevation and myopathy in a BAT-dependent manner. **(A)** *Irf4* mRNA levels in BAT from mice at RT and 30ºC. (**B**) Western blot analysis of isolated mitochondria from vastus of WT mice at RT and 30°C. (**C**) Western blot analysis of mTOR signaling pathway in vastus of WT mice at RT and 30°C. Activity was quantified using a phosphorimager (n=4, \**p* < 0.05 vs. RT mice). (**D**) Q-PCR analysis of vastus gene expression (n=5-6, \**p* < 0.05). **(E)** Q-PCR analysis of vastus ribosomal gene expression (n=5-6, \**p* < 0.05). **(F)** Body weight (BW) of iBATx and sham-operated mice after surgery. **(G)** Q-PCR analysis of vastus gene expression before and after iBATx and exposure to 30°C (n=5-6, \**p* < 0.05). **(H)** Q-PCR analysis of vastus ribosomal gene expression before and after iBATx and exposure to 30°C (n=5-6, \**p* < 0.05). **(I)** Western blot analysis of mTOR signaling pathway in vastus from mice before and after iBATx and exposure to 30°C. Activity was quantified using a phosphorimager. (n=4, \**p* < 0.05 vs. Flox mice. #*p* < 0.05 vs. iBATx mice.)

**Supplemental Table S1:**
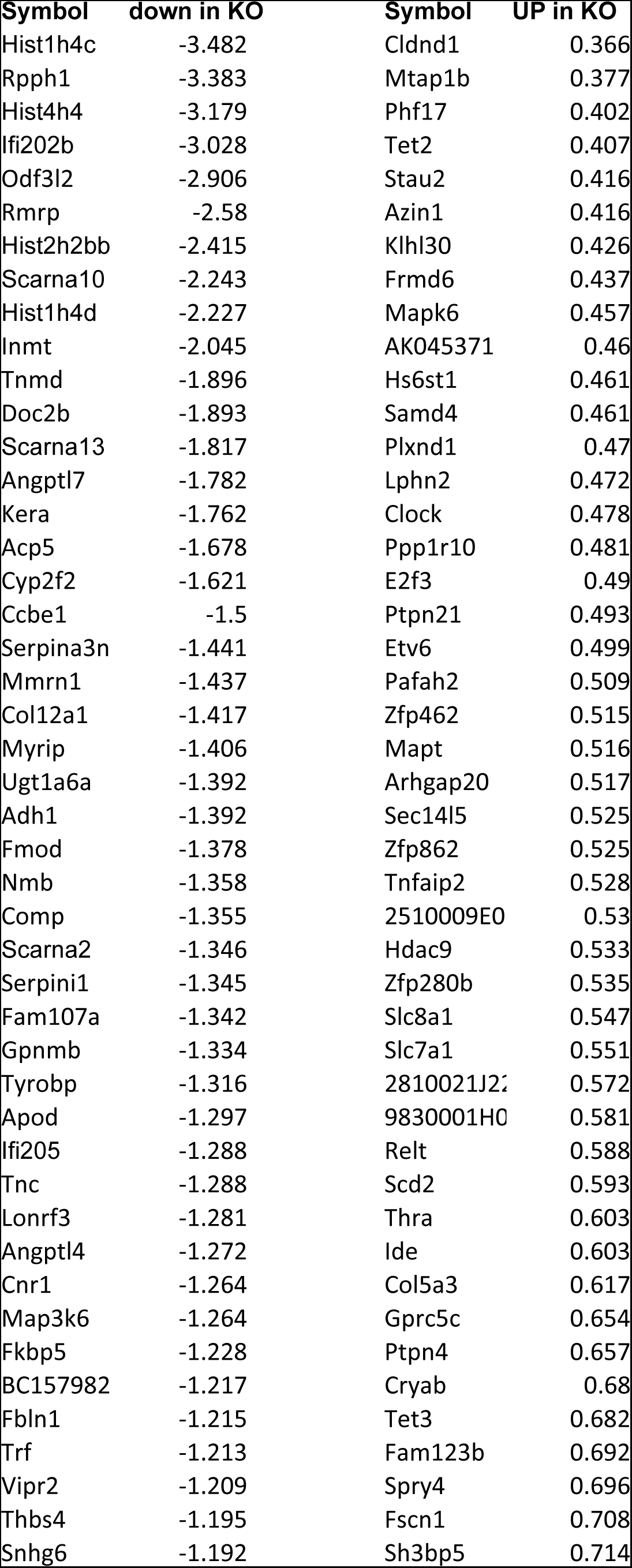
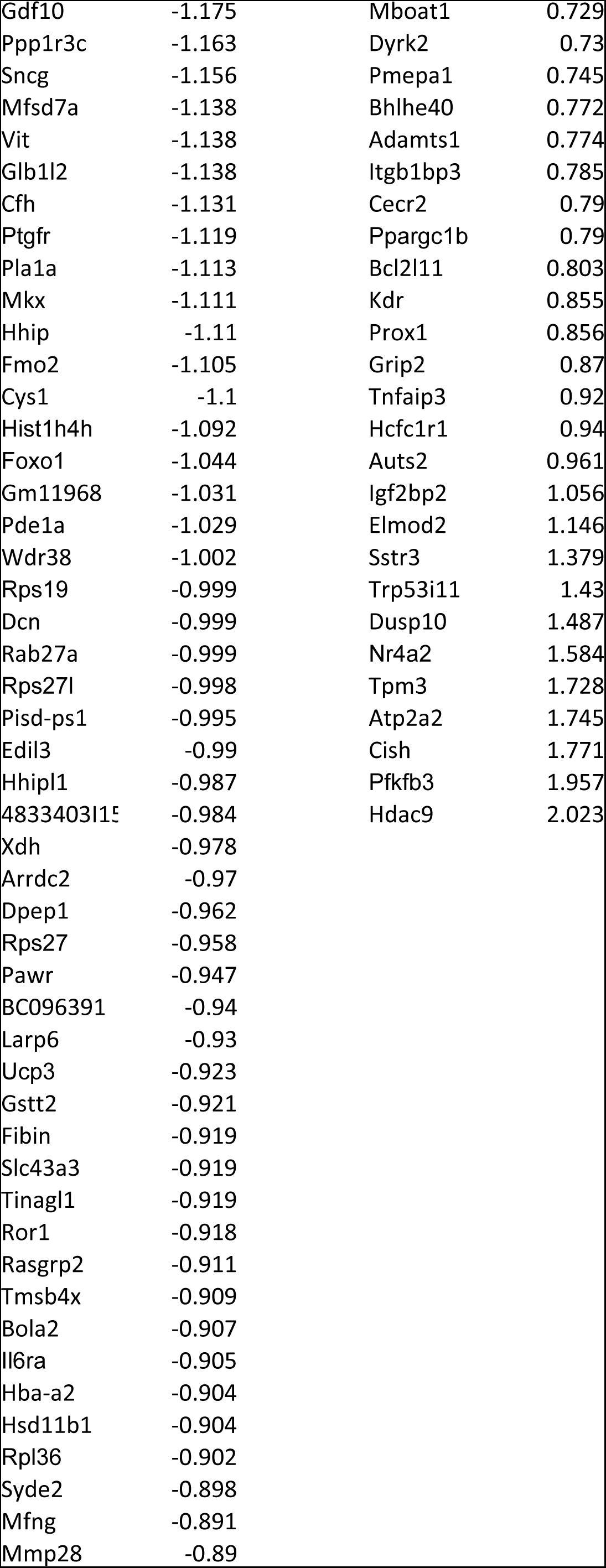
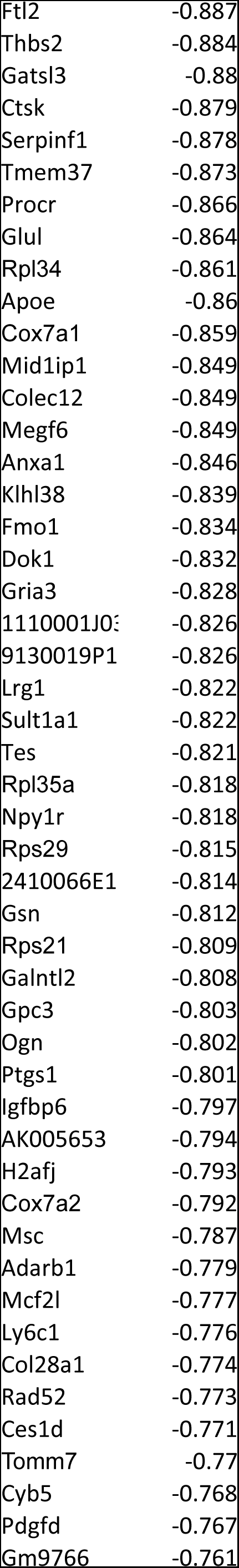
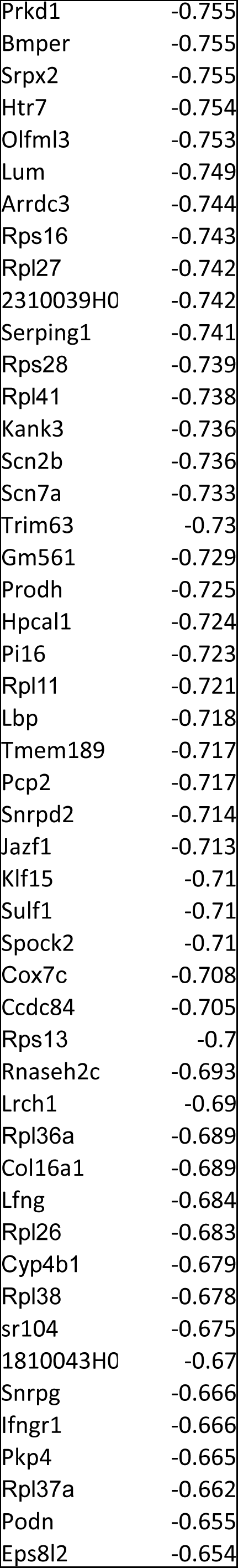
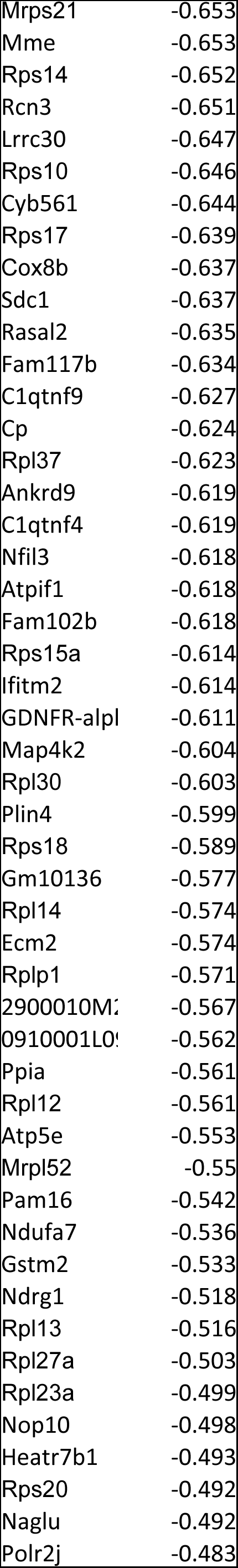
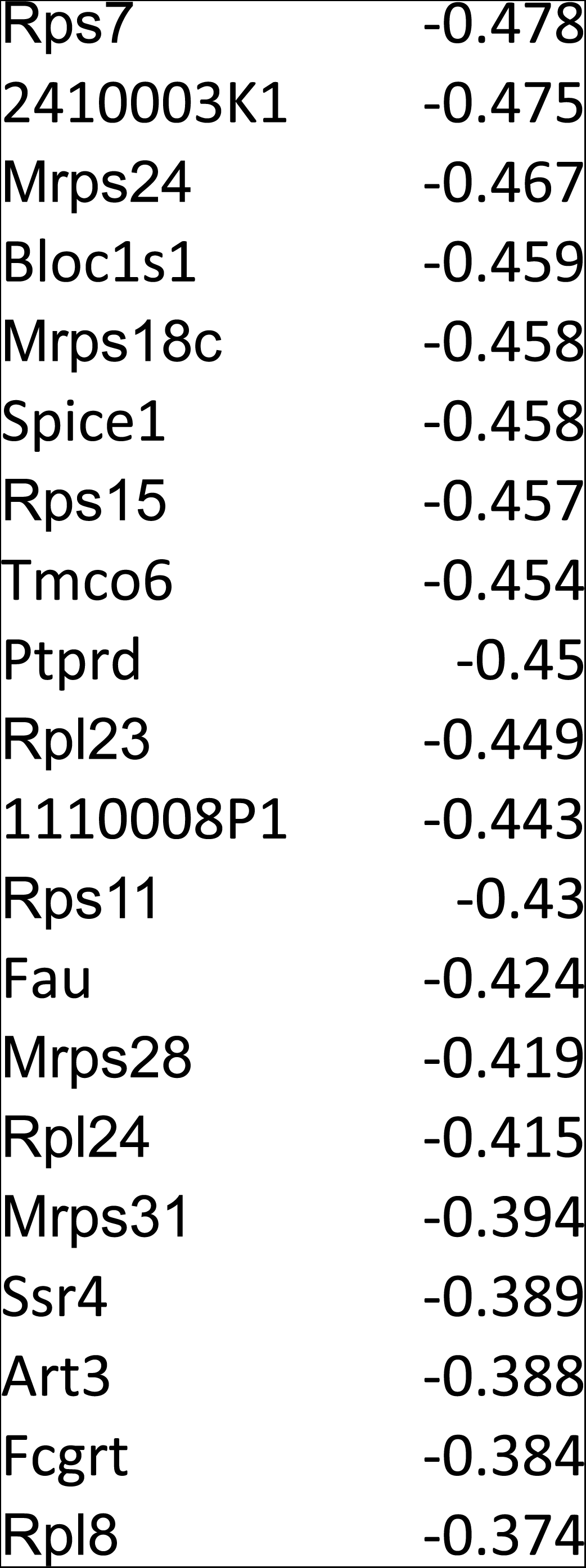
Up- and down-regulated genes in vastus lateralis of BATI4KO vs. control mice. Data are presented as log2 fold change.

**Supplemental Table S2:**
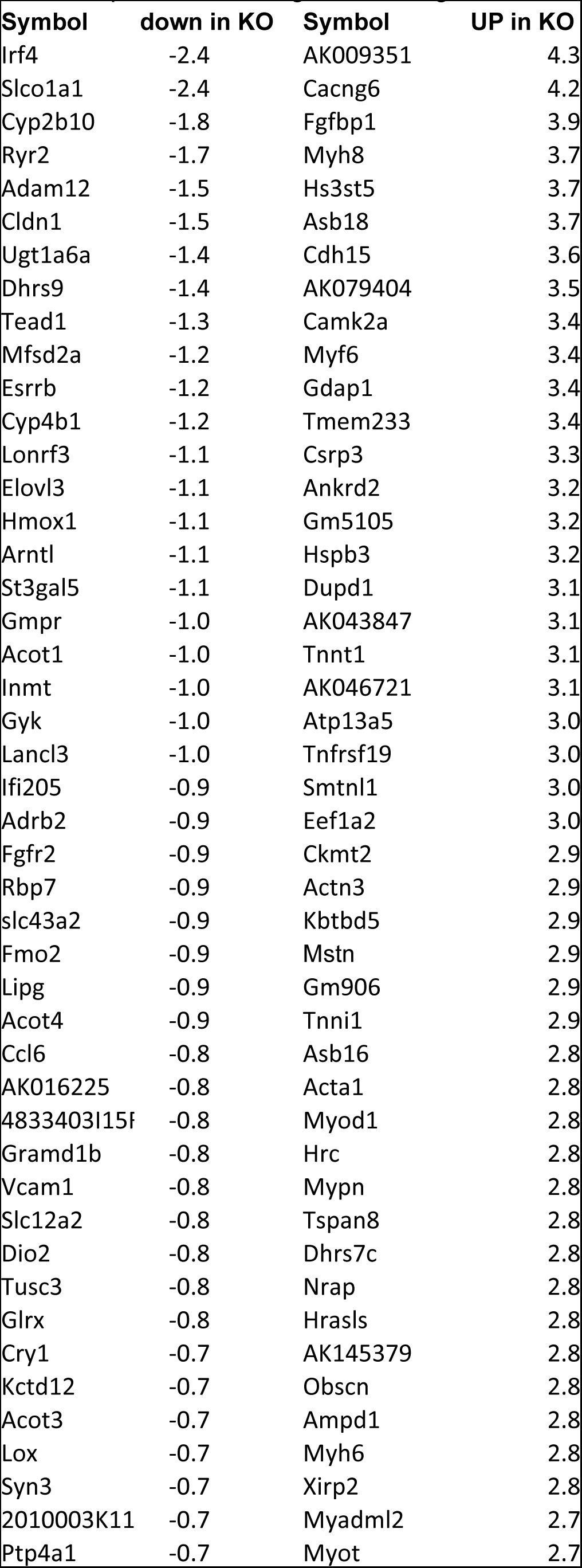
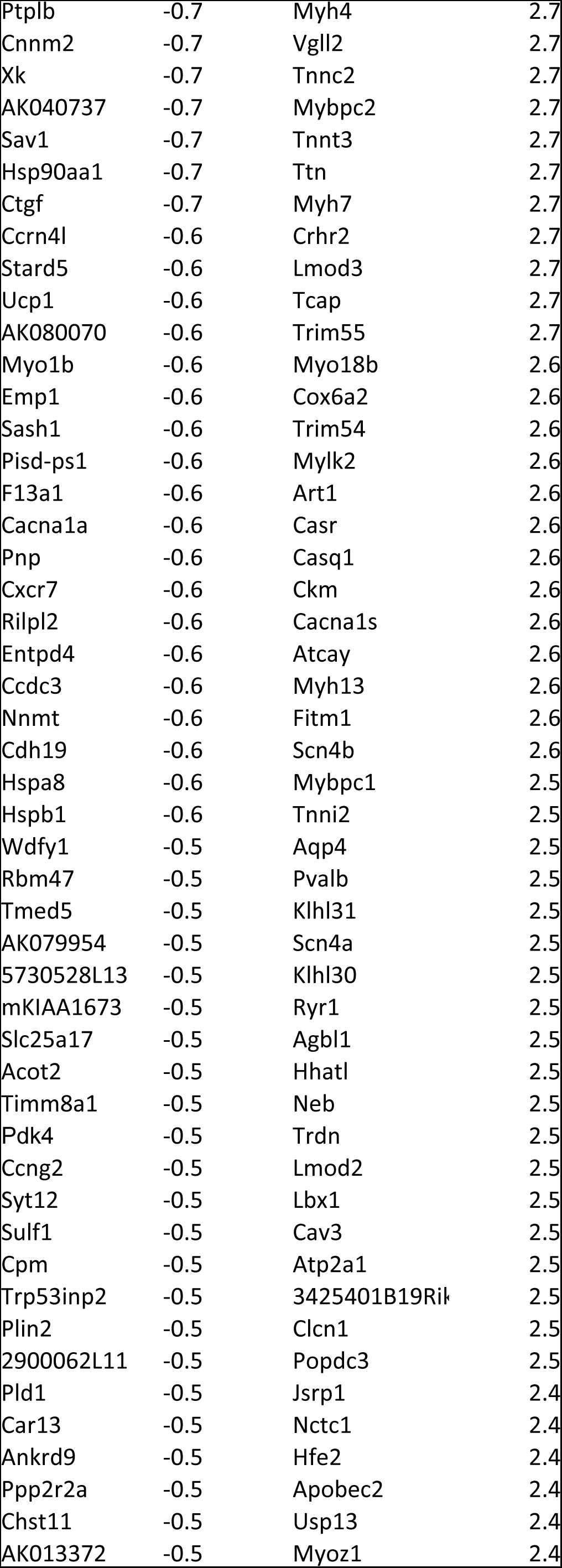
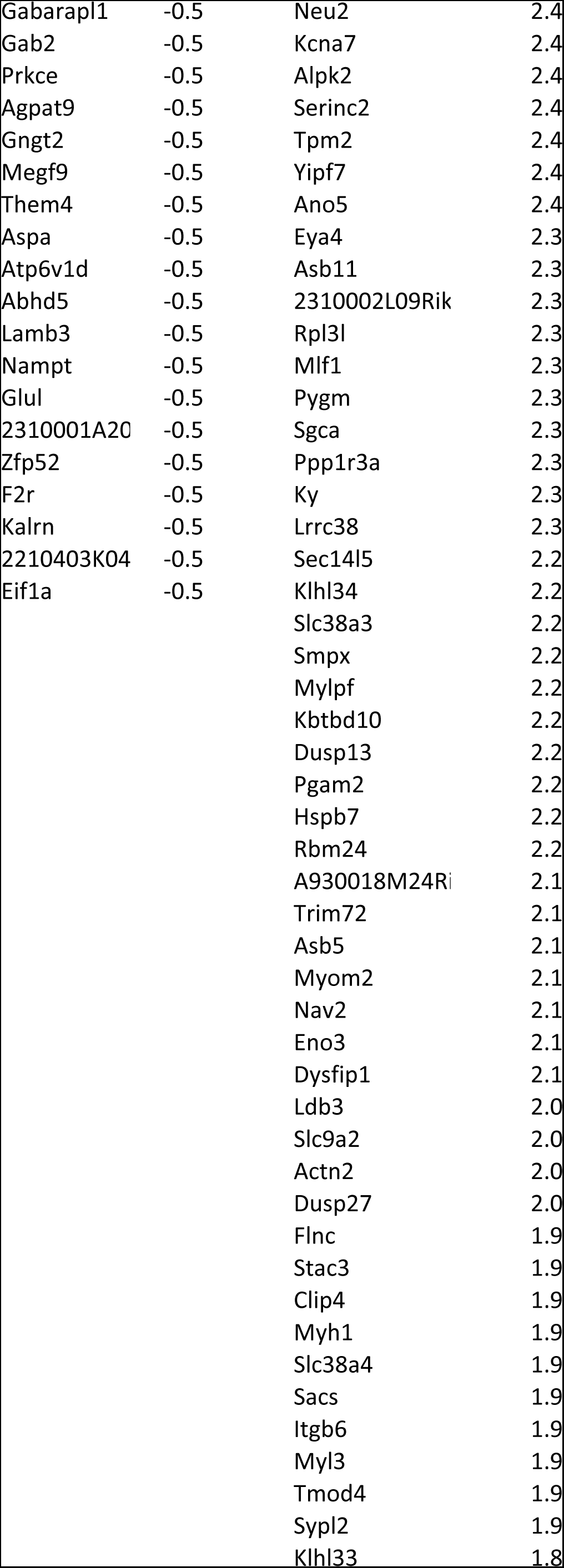
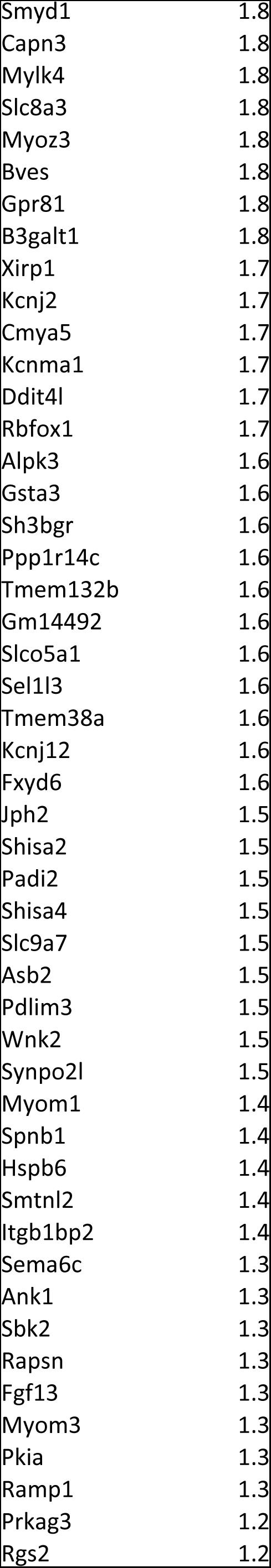
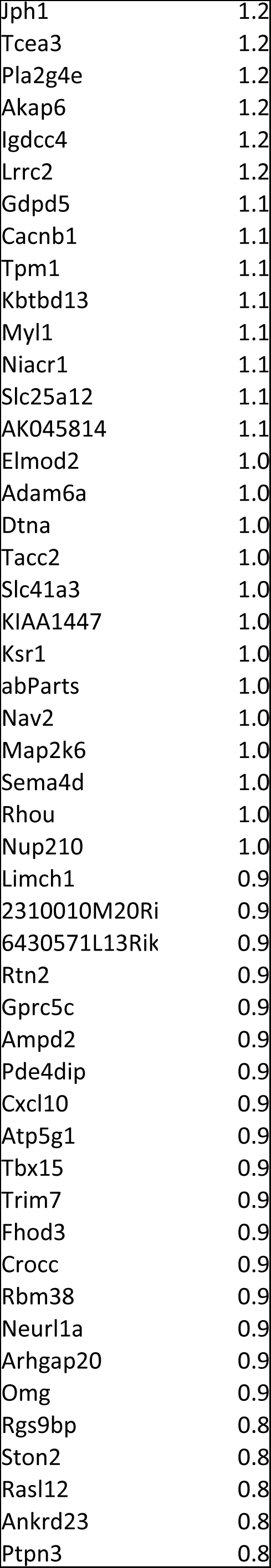
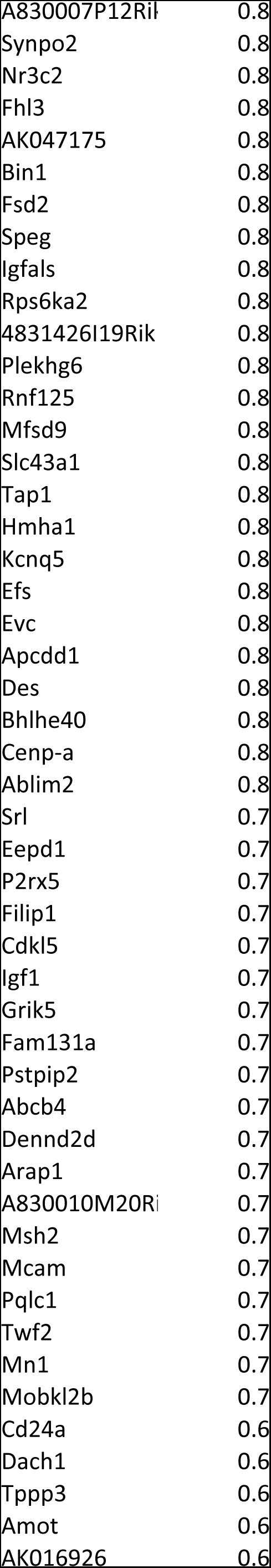
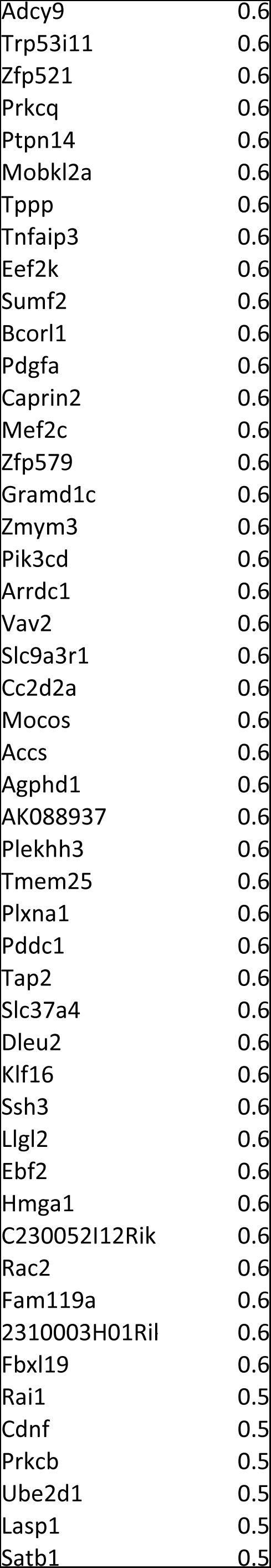
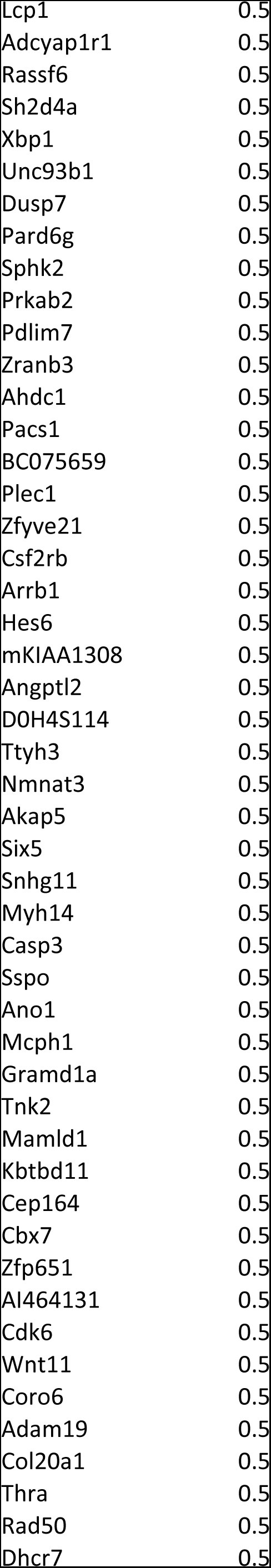
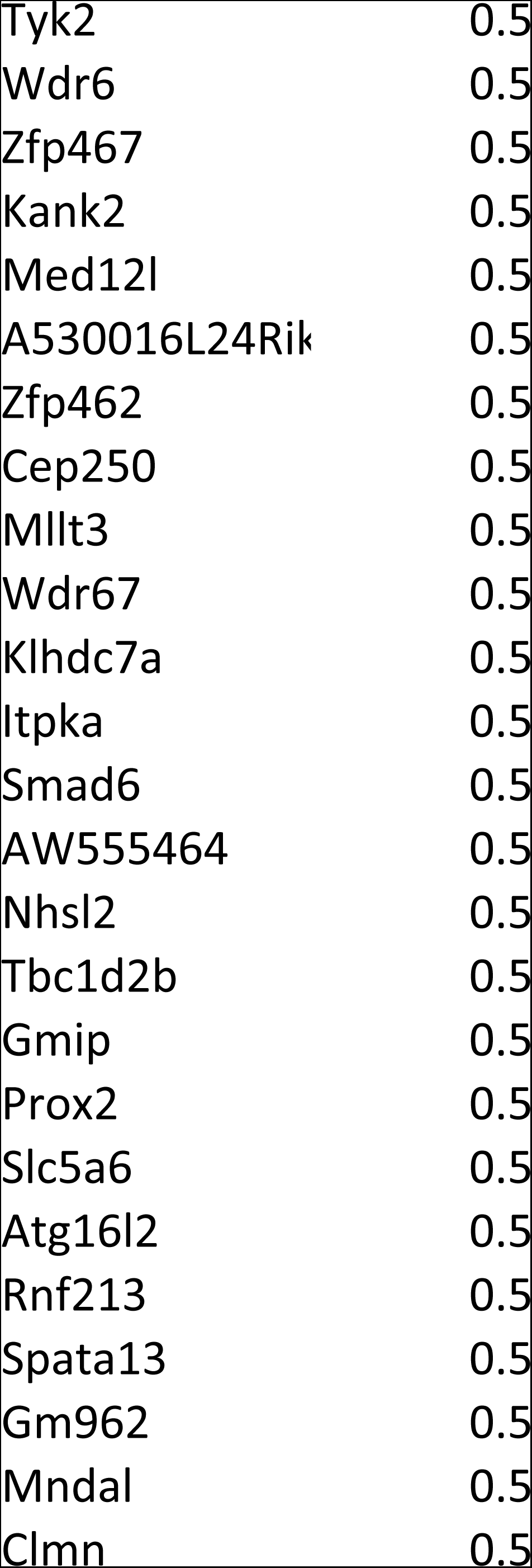
Up- and down-regulated genes in brown adipose tissue of BATI4KO vs. control mice. Data are presented as log2 fold change.

